# De Novo Mutational Signature Discovery in Tumor Genomes using SparseSignatures

**DOI:** 10.1101/384834

**Authors:** Avantika Lal, Keli Liu, Robert Tibshirani, Arend Sidow, Daniele Ramazzotti

## Abstract

Cancer is the result of mutagenic processes that can be inferred from tumor genomes by analyzing rate spectra of point mutations, or “mutational signatures”. Here we present SparseSignatures, a novel framework to extract signatures from somatic point mutation data. Our approach incorporates a user-specified background signature, employs regularization to reduce noise in non-background signatures, uses cross-validation to identify the number of signatures, and is scalable to large datasets. We show that SparseSignatures outperforms current state-of-the-art methods on simulated data using a variety of standard metrics. We then apply SparseSignatures to whole genome sequences of pancreatic and breast tumors, discovering well-differentiated signatures that are linked to known mutagenic mechanisms and are strongly associated with patient clinical features.

**Authors Summary:** Cancer is a genetic disease, occurring as a result of mutagenic processes causing DNA somatic mutations in genes controlling cellular growth and division. These somatic mutations arise from processes such as defective DNA repair and environmental mutagens, which massively increase the rate of somatic variants. As a result, due to the specificity of molecular lesions caused by such processes, and the specific repair mechanisms deployed by the cell to mitigate the damage, mutagenic processes generate characteristic point mutation rate spectra which are called mutational signatures. These signatures can indicate which mutagenic processes are active in a tumor, reveal biological differences between cancer subtypes, and may be useful markers for therapeutic response. Here, we develop SparseSignatures, a novel framework for mutational signature discovery capable of both identifying the active signatures in a dataset of point mutations and calculating their exposure values, i.e., the number of mutations originating from each signature in each patient. We show that our approach outperforms current state-of-the-art methods on simulated data using a variety of standard metrics and then apply SparseSignatures to whole genome sequences of pancreatic and breast tumors, discovering well-differentiated signatures that are linked to known mutagenic mechanisms.

## Introduction

Cancer is caused by somatic mutations in genes that control cellular growth and division [1]. The chance of developing cancer is massively elevated if mutagenic processes (e.g., defective DNA repair, environmental mutagens) increase the rate of somatic mutations. Due to the specificity of molecular lesions caused by such processes, and the specific repair mechanisms deployed by the cell to mitigate the damage, mutagenic processes generate characteristic point mutation rate spectra (‘signatures’) [2]. These signatures can indicate which mutagenic processes are active in a tumor, reveal biological differences between cancer subtypes, and may be useful markers for therapeutic response [3].

Signatures are discovered by identifying common patterns across tumors based on counts of mutations and their sequence context. The original signature discovery method was based on Non-Negative Matrix Factorization (NMF) [4]. While other approaches have been considered [5,6], NMF-based methods are by far the most widely used [7–9] and have resulted in a commonly used catalog of 30 signatures across human cancers [10], available in the COSMIC version 2 database (https://cancer.sanger.ac.uk/cosmic/signatures_v2). A recent study [11] using two NMF-based methods presented higher numbers (49 and 60) of putative signatures, which has now been incorporated into version 3 of the COSMIC database (https://cancer.sanger.ac.uk/cosmic/signatures).

While some reported signatures have been associated with mutagenic processes [9,12,13], careful examination reveals that several reported signatures are highly similar, suggesting overfitting rather than distinct mutagenic processes. In addition, there are several ‘flat’ signatures of uncertain origin (non-specific signatures that include mutations of all types and sequence contexts), and many signatures appear to be distorted by low levels of background noise. As an example, one may consider SBS40 in COSMIC version 3, whose etiology is unclear and which has many features in common with SBS5 [11]. Another example is represented by the four similar signatures in COSMIC version 2 that are attributed to defective DNA mismatch repair (signatures 6, 15, 20, and 26), which share common features and are not clearly separated. Such uncertainty complicates the task of understanding which signatures are active in different patients. These observations are consistent with critical weaknesses in current signature discovery studies:

1. State-of-the-art NMF-based methods aim to minimize the residual error after fitting the dataset with the discovered signatures [4,5], in an effort to fit the dataset perfectly. Consequently they may overfit by including stochastic noise in the dataset as part of the signatures, or by producing multiple similar signatures for the same underlying process. This problem is exacerbated by the relatively low number of samples (hundreds or thousands) available to most mutational signature discovery studies. LASSO regularization has been shown to improve estimation in high dimensional problems when the sample size is small relative to the number of parameters [14]. A method that applies LASSO regularization on the signatures would help alleviate the aforementioned drawbacks by favoring well-differentiated signatures with low background noise, in addition to minimizing residual error. Variants of NMF that incorporate regularization are available and have been used in other domains [15,16], the NNLM R package on CRAN at https://cran.r-project.org/web/packages/NNLM/index.html), and a few recent studies [17,18] have attempted to apply these methods to signature discovery.
2. Many independent studies have found a highly dense (‘flat’) signature (SBS5) to be abundant in diverse settings, including all human cancer types profiled in COSMIC [10] and PCAWG [11], numerous cancer cell lines [19] non-cancer somatic tissues [20,21], adult stem cell-derived organoids [22], 1000 Genomes Project SNPs from different human populations [23] and germline *de novo* mutations [24]. We discuss the etiology of this signature later in the paper. Mathematically, the high density of this signature renders it difficult to accurately extract *de novo*, especially under the conditions of low mutational rates, few samples, and multiple flat signatures, all of which are common. When not fitted accurately, this signature may contaminate other signatures leading to inaccurate estimation and assignment of signatures. Considering these potential issues and the ubiquity of SBS5, a recent prominent pan-cancer study across all PCAWG samples deliberately assigned SBS5 to be present in all samples [11]. However, SBS5 was not fixed as part of the signature discovery method itself but was assigned to samples afterward, which does not resolve the problem of contamination of other signatures. This procedure can be improved using matrix factorization methods that allow for fixing some elements of the solution [16,25,26], i.e., fixing one or more signatures as a constant.
3. State-of-the-art NMF-based methods require the number of signatures as an input parameter but lack a principled basis for its selection. Discovering more signatures will always tend to reduce the residual error, i.e., fit the observed data better. However, the goal of signature discovery is not only to fit the data as well as possible, but also to identify signatures that truly reflect separate biological processes. Currently, standard ways to choose the number of signatures are: (1) choosing a number such that more signatures would not significantly reduce residual error [5]; (2) choosing a number based on both minimizing residual error and maximizing reproducibility of signatures [4]; (3) calling signatures hierarchically on subsets of samples, adding more signatures in order to fit every sample [9]. The first two practices are ambiguous, while the third selects as many signatures as needed to improve fitting of the data, with little constraint to prevent overfitting. Overfitting can lead to many spurious signatures that actually represent noise, making it difficult to reliably attribute mutations in a sample to any one signature, leading to misinterpretation of the results and misleading conclusions. However, successful methods have been developed to choose the number of factors in NMF, including missing values imputation (the NNLM R package) and cross-validation [27]. A recent signature discovery method, SignatureAnalyzer, uses automatic relevance determination, which starts with a high number of signatures and attempts to eliminate signatures of low relevance [28].

To overcome these drawbacks, we developed SparseSignatures (Figure 1A), a novel framework for mutational signature discovery. Like other NMF-based methods, SparseSignatures both identifies the signatures in a dataset of point mutations and calculates their exposure values (the number of mutations originating from each signature) in each patient.

**Figure 1.**
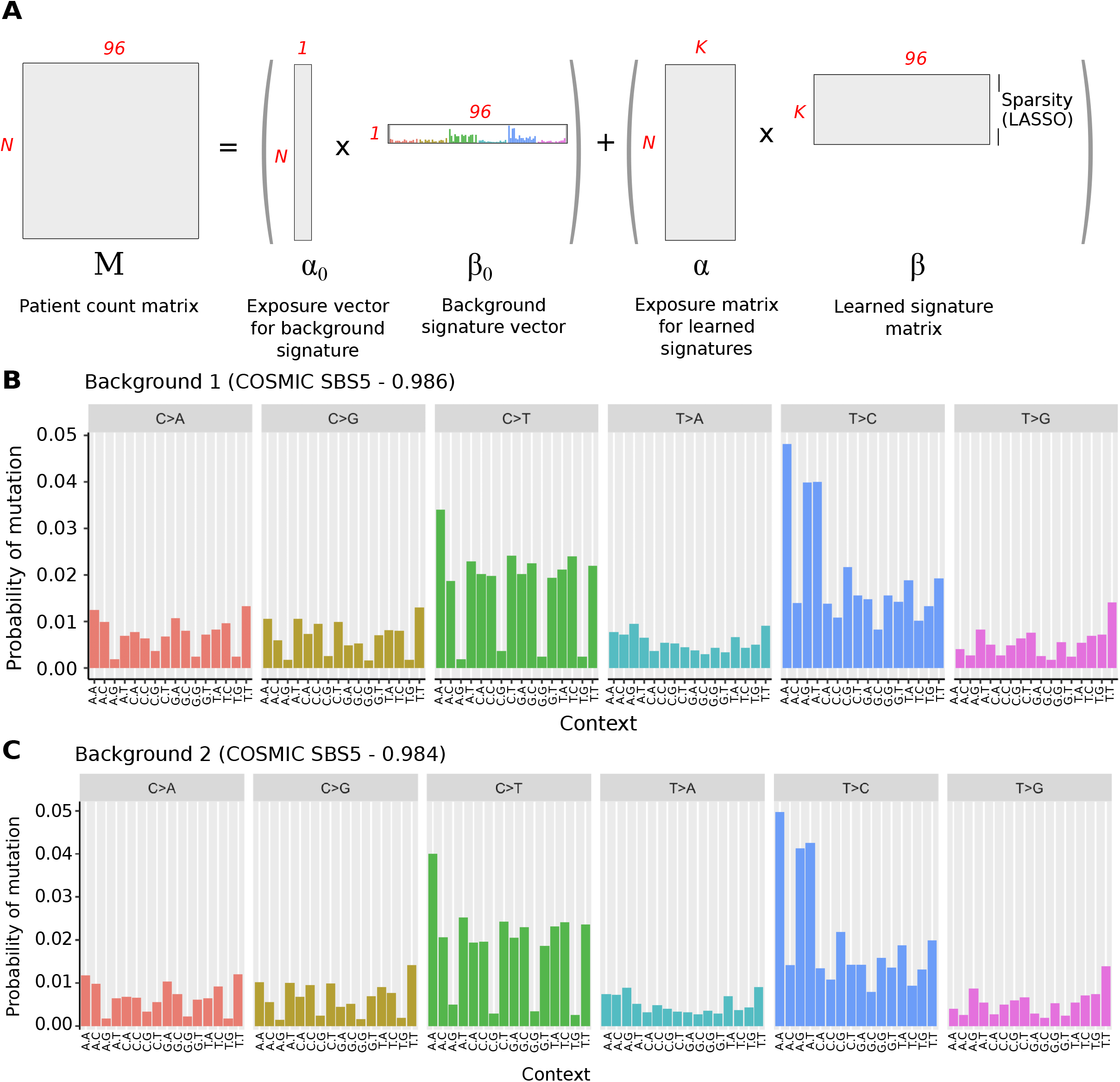
A) Schematic of the SparseSignatures method. N represents the number of tumors in the dataset, K the number of signatures. B) Background signature derived from COSMIC SBS5. C) Background signature derived from the human germline mutation spectrum. Vertical bars represent the probability of mutation in each of 96 categories. These are based on 6 possible mutation types (upper gray labels) and 16 possible combinations of 5’ and 3’ flanking bases (x-axis labels). Source data are provided in Supplementary Table 1.

## Results

### The SparseSignatures Algorithm

SparseSignatures is implemented in R and is available as a Bioconductor package at https://bioconductor.org/packages/release/bioc/html/SparseSignatures.html. Noteworthy innovations are:

**Table 1.** Signatures (including background) discovered by SparseSignatures in real cancer data, and their proposed etiology.

| Signature | Proposed etiology | Basis for proposed etiology |
| --- | --- | --- |
| Pancreatic cancer (147 whole genomes) |  |  |
| PC-SS1 | Cytosine methylation / deamination | Experimental evidence [13] |
| PC-SS2 | APOBEC dysregulation | Experimental evidence [13] |
| PC-SS3 | Defective homologous recombination-based DNA damage repair | Hypothesised [11] |
| Background | DNA replication error | Experimental evidence [24] |
| PC-SS4 | APOBEC dysregulation | Experimental evidence [13] |
| PC-SS5 | Oxidative damage | Hypothesized [40] |
| PC-SS6 | Damage by reactive oxygen species | Hypothesised [11] |
| PC-SS7 | Defective DNA mismatch repair | Hypothesised [11] |
| PC-SS8 | Possible sequencing artefact | Hypothesised [11] |
| Breast cancer (560 whole genomes) |  |  |
| BRCA-SS1 | Cytosine methylation / deamination | Experimental evidence [13] |
| BRCA-SS2 | APOBEC dysregulation | Experimental evidence [13] |
| BRCA-SS3 | Defective homologous recombination-based DNA damage repair | Hypothesised [11] |
| Background | DNA replication error | Experimental evidence [24] |
| BRCA-SS4 | APOBEC dysregulation | Experimental evidence [13] |
| BRCA-SS5 | Oxidative damage | Hypothesized [40] |
| BRCA-SS6 | Damage by reactive oxygen species | Hypothesised [11] |
| BRCA-SS7 | Defective DNA mismatch repair | Hypothesised [11] |

1. It allows the user to incorporate an explicit background model of their choice by specifying a fixed ‘background’ signature. Two preset background signatures are provided. One (Figure 1B, Supplementary Table 1) was derived from the SBS5 signature in COSMIC, which has been found in all studied cancer types as well as normal somatic tissue [10,20] and has been considered a natural background signature [29]. The other (Figure 1C, Supplementary Table 1) was derived from the human germline mutation spectrum [24], and validated in normal tissue samples (Supplementary Methods). For both of these, we made an empirical adjustment to CpG > TpG mutation rates (see Methods). This is because CpG > TpG mutations are frequently caused by cytosine deamination at sites of CpG methylation. Since the extent of CpG methylation can vary greatly in cancer cells, the rate of such mutations is not perfectly correlated with replication rates in tumors. The cosine similarity between these two background signatures is 0.998 and they provide almost identical results. SparseSignatures fixes the background signature and then discovers additional signatures representing cancer-specific mutagenic processes (including, usually, deamination of methylated cytosines). Moreover, users can choose to use no background signature, or to provide a background signature of their choice.
2. It uses LASSO regularization [14] to reduce noise in the signatures, except for the fixed background signature (if provided). The extent of regularization is controlled by a learned parameter, λ, for the entire signature matrix. We note that if the underlying signatures are very different in sparsity, this could result in a few individual signatures being too sparse or too dense if the value of λ is not ideal for them. However, we aim to improve the overall solution, and so our method chooses the best overall value of λ based on the complete dataset. It is also capable of choosing λ=0 (no LASSO penalty) if regularization does not in fact improve the solution.
3. It implements repeated bi-cross-validation [30] to select the best values for both the regularization parameter (λ) and the number of signatures (K). A randomly chosen subset of data points is held out and signatures are discovered based on the rest of the data. The values of the held-out data points are predicted based on the discovered signatures and their fitted exposure values in each patient, and the mean squared error of the predictions is calculated. This procedure is performed for different values of K and λ, and the values that minimize the error in predicting held-out data points are chosen. The goal is to avoid overfitting, by ensuring that the discovered signatures not only fit the data used for discovery but also predict unseen values with high accuracy. In contrast to several previous methods, this provides a clear, unambiguous metric to choose the number of signatures.

### SparseSignatures accurately deciphers signatures in simulated data

We compared SparseSignatures to two existing NMF-based methods for signature discovery, SigProfiler [4,11] and SignatureAnalyzer [28]. SigProfiler and SignatureAnalyzer were the basis for a recent pan-cancer study [11] resulting in 49 and 60 putative signatures. We also included signeR [31], a Bayesian approach. In *Simulation 1*, we generated 50 simulated datasets of 116 patients each with 4 underlying mutational signatures, based on curated WGS data from a cohort of Prostate cancer patients (see Methods). The underlying mutational signatures included a dense signature (COSMIC SBS3) as well as relatively sparse signatures (COSMIC SBS1, SBS18). We applied all four methods for signature discovery to this simulated dataset.

On this simulated data, SparseSignatures is most effective at discovering the correct number of signatures (Figure 2A, Supplementary Table 2). SignatureAnalyzer consistently overfits the data, i.e., it overestimates the number of signatures and discovers an excessive number of sparse signatures that fit the data well but do not represent the actual underlying processes.

**Table 2.** Comparison of signatures predicted by four signature discovery methods on real tumor sequencing data. Sparsity is measured as the fraction of cells in the signature matrix with value < 10^−4^. Cross-signature similarity is measured as the mean cosine similarity between all pairs of predicted signatures. Background contamination is measured as the mean cosine similarity between the background signature and all non-background predicted signatures. Median per-patient correlation is measured as the median Pearson’s correlation coefficient between the observed mutation spectrum and the predicted mutation spectrum for each patient, indicating how well each method fits the observed mutations in individual patients.

| Source | Number of signatures | MSE | Sparsity (signatures) | Cross-signature similarity | Background contamination | Median per-patient correlation |
| --- | --- | --- | --- | --- | --- | --- |
| Pancreatic cancer (147 whole genomes) |  |  |  |  |  |  |
| SparseSignatures | 9 | 1189.521 | 0.317 | 0.193 | 0.373 | 0.9916 |
| SigProfiler | 8 | 52564.38 | 0.0762 | 0.384 | 0.514 | 0.9910 |
| SignatureAnalyzer | 8 | 52606.28 | 0.245 | 0.240 | 0.447 | 0.9911 |
| signeR | 8 | 52733.45 | 0.003 | 0.320 | 0.502 | 0.9908 |
| Breast cancer (560 whole genomes) |  |  |  |  |  |  |
| SparseSignatures | 8 | 1515.373 | 0.154 | 0.254 | 0.387 | 0.9762 |
| SigProfiler | 12 | 1034.204 | 0.060 | 0.275 | 0.452 | 0.9360 |
| SignatureAnalyzer | 12 | 72191.45 | 0.449 | 0.158 | 0.300 | 0.9805 |
| signeR | 7 | 72390.55 | 0.115 | 0.317 | 0.458 | 0.9735 |

**Figure 2.**
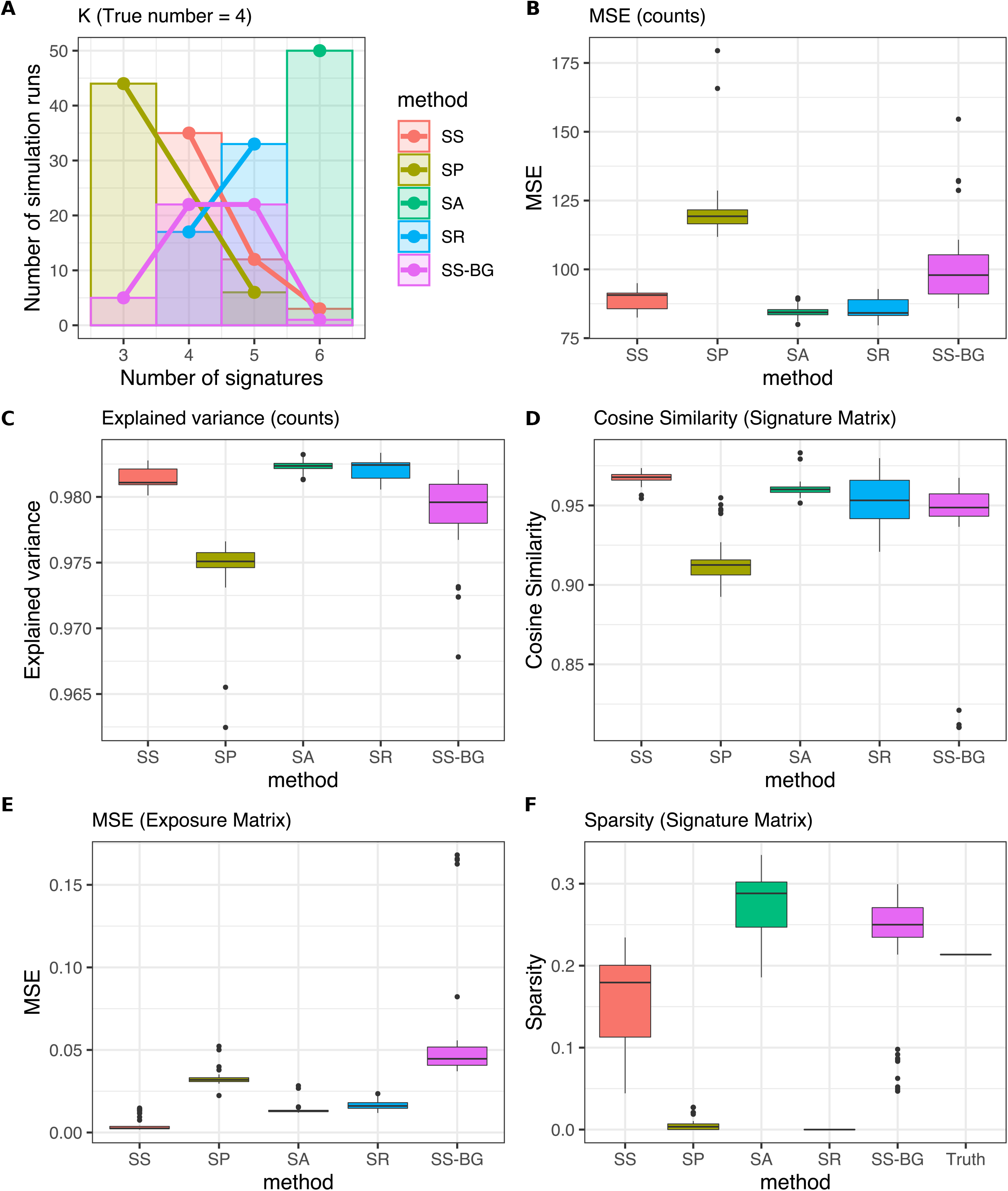
Comparison between SparseSignatures and other methods on simulated data. A) Bar and line plot showing, for each method, the number of simulations in which it selected each value of K (number of signatures). The x-axis shows values of K and the y-axis shows the number of times each value was selected. Each method was run on 50 simulated datasets. In all cases, the correct value of K was 4. B) Box plots showing the residual error for the solutions produced by each method, over 50 simulations. Residual error was measured as the mean squared error (MSE) in reconstructing the original count matrix. C) Box plots showing the fraction of variance in the count matrix explained by the solutions produced by each method, over 50 simulations. D) Box plots showing the cosine similarity in reconstructing the 3 non-background input signatures, over 50 simulations. E) Box plots showing the mean squared error in reconstructing the exposure values for the 3 non-background input signatures, over 50 simulations. F) Box plots showing the sparsity of the signatures produced by each method, over 50 simulations. Sparsity was measured as the fraction of cells in the signature matrix whose value is <10^−3^. SS: SparseSignatures. SP: SigProfiler. SA: SignatureAnalyzer. SR: signeR. SS-BG: SparseSignatures without fixed background. Source data are provided in Supplementary Table 2.

When comparing the overall residual error obtained by the four methods, SignatureAnalyzer fits the input matrix with the least residual error (Figure 2B-C, Supplementary Table 2). However, this is the result of overfitting as the method infers too many signatures. To provide a clearer measure, we assessed how well each method deciphers the input signatures by matching each of the input signatures to the most similar signature produced by the method, and assessing the cosine similarity between these pairs of signatures. We did not include the background signature in this comparison. Compared to all other methods, SparseSignatures reconstructs the input signatures more accurately (Figure 2D, Supplementary Table 2). Supplementary Figures 1 and 2, along with Supplementary Table 3, show the simulated patient counts, original signatures, and signatures predicted by each method, for one of the 50 simulated datasets. We also compared the original exposure values for each input signature to the exposure values produced by the method for the closest deciphered signature, and found that SparseSignatures shows the lowest error in reconstructing the original exposure values (Figure 2E, Supplementary Table 2).

Appropriate regularization of the signatures based on a learned parameter (λ) is one reason for the higher accuracy of our approach. The sparsity of signatures deciphered by SparseSignatures closely matches that of the input signatures (Figure 2F, Supplementary Table 2). In comparison, SigProfiler and signeR tend to discover signatures with the addition of considerable noise, while SignatureAnalyzer produces excessively sparse signatures. We also demonstrated that the superior performance of SparseSignatures depends upon the inclusion of a fixed background signature; if this signature is not fixed, SparseSignatures is unable to accurately reconstruct it and other signatures, and the performance of our method is reduced across all metrics (Fig. 2A-F, Supplementary Table 2).

Finally, while SparseSignatures is the most accurate method at discovering the correct number of signatures, we also compared the performance of all three methods if the correct number of signatures is already known. When all four methods were given the correct number of signatures, SparseSignatures was still the most accurate at reconstructing the input signatures and exposures (Supplementary Figure 3, Supplementary Table 4).

**Figure 3.**
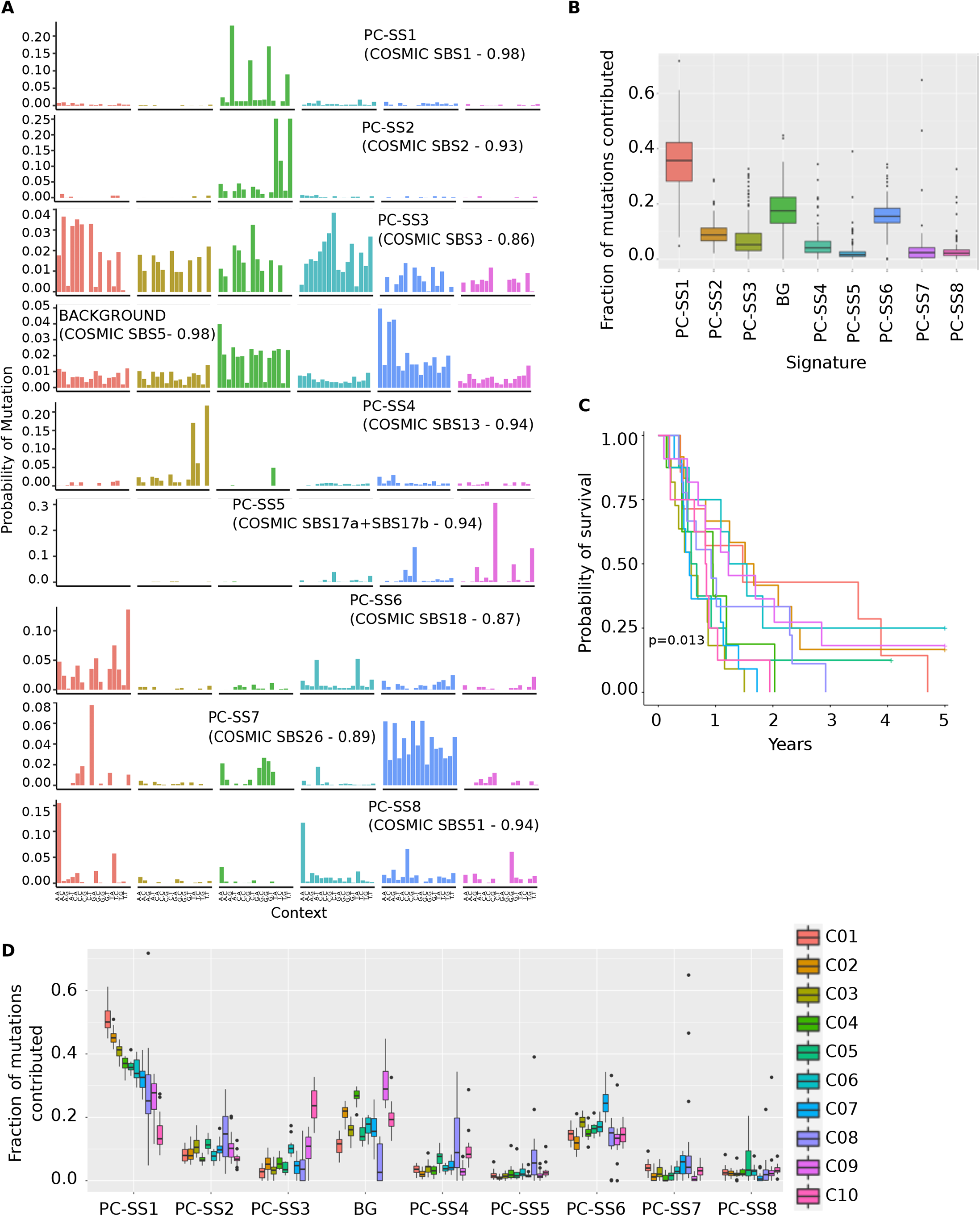
A) The 9 mutational signatures obtained by applying SparseSignatures to a dataset of 147 pancreatic tumors. We report the number and correlation of the most similar (correlation higher than 0.70) corresponding signature from COSMIC (https://cancer.sanger.ac.uk/cosmic/signatures). Source data are provided in Supplementary Table 13. B) Boxplots showing fitted values for exposure to each of the 9 signatures obtained by SparseSignatures for the 147 pancreatic tumors. Boxplots represent the fraction of mutations per tumor (on the y-axis) contributed by the given signature (on the x-axis). Source data are provided in Supplementary Table 14. C) Relapse-free survival analysis of patients belonging to the 10 clusters. D) Clustering of patients based on their exposure values. Boxplots show the fraction of mutations per tumor contributed by each signature (x-axis) to each of 10 clusters. Source data are provided in Supplementary Table 19.

To provide additional validation of the robust performance of SparseSignatures, we performed three additional simulation experiments with different types of underlying signatures.

1. *Simulation 2*: We generated 50 simulated datasets of 116 patients each with 4 underlying mutational signatures as in Simulation 1, but including a wider range of noise.
2. *Simulation 3*: We generated 50 simulated datasets, each of which used 4 randomly selected signatures from the COSMIC version 3 database.
3. *Simulation 4*: We generated 50 simulated datasets, each of which used 4 randomly selected signatures from the COSMIC version 3 database, limited to relatively dense signatures where >75% of the 96 mutation types contribute to the signature.
4. *Simulation 5*: We generated 50 simulated datasets, each of which used 4 randomly selected signatures from the COSMIC version 3 database, limited to relatively sparse signatures where <50% of the 96 mutation types contribute to the signature.
5. *Simulation 6*: We generated 50 simulated datasets, each of which contained 100 simulated patients with 8 underlying mutational signatures selected from the COSMIC version 3 database.

In all these additional simulations, we obtained similar results (Supplementary Figures 4-8, Supplementary Tables 5-9). SignatureAnalyzer performs poorly at discovering the number of signatures; SparseSignatures, SigProfiler, and signeR all perform better, frequently identifying the correct number of signatures or coming close (Supplementary Figures 4A,5A, 6A, 7A, 8A). However, SparseSignatures is more accurate than SigProfiler at reconstructing both the input signatures (Supplementary Figures 4D, 5D, 6D, 7D, 8D) and exposures (Supplementary Figures 4E, 5E, 6E, 7E, 8E). While signeR also performs well at reconstructing the signature matrix, SparseSignatures consistently exceeds the performance of signeR at reconstructing the exposure matrix (Supplementary Figures 4E, 5E, 6E, 7E, 8E). It is particularly notable that across all simulations, both SigProfiler and signeR recover signatures with considerable background noise. This is in clear contrast to SparseSignatures, which, due to the combination of regularization and fixing the background signature, minimizes background noise and recovers signatures of the correct sparsity (Supplementary Figures 4F, 5F, 6F, 7F, 8F). Overall, SparseSignatures exceeds the performance of all the other methods. This shows the robust performance of SparseSignatures and its ability to accurately reconstruct input signatures and exposures from datasets with different characteristics.

**Figure 4.**
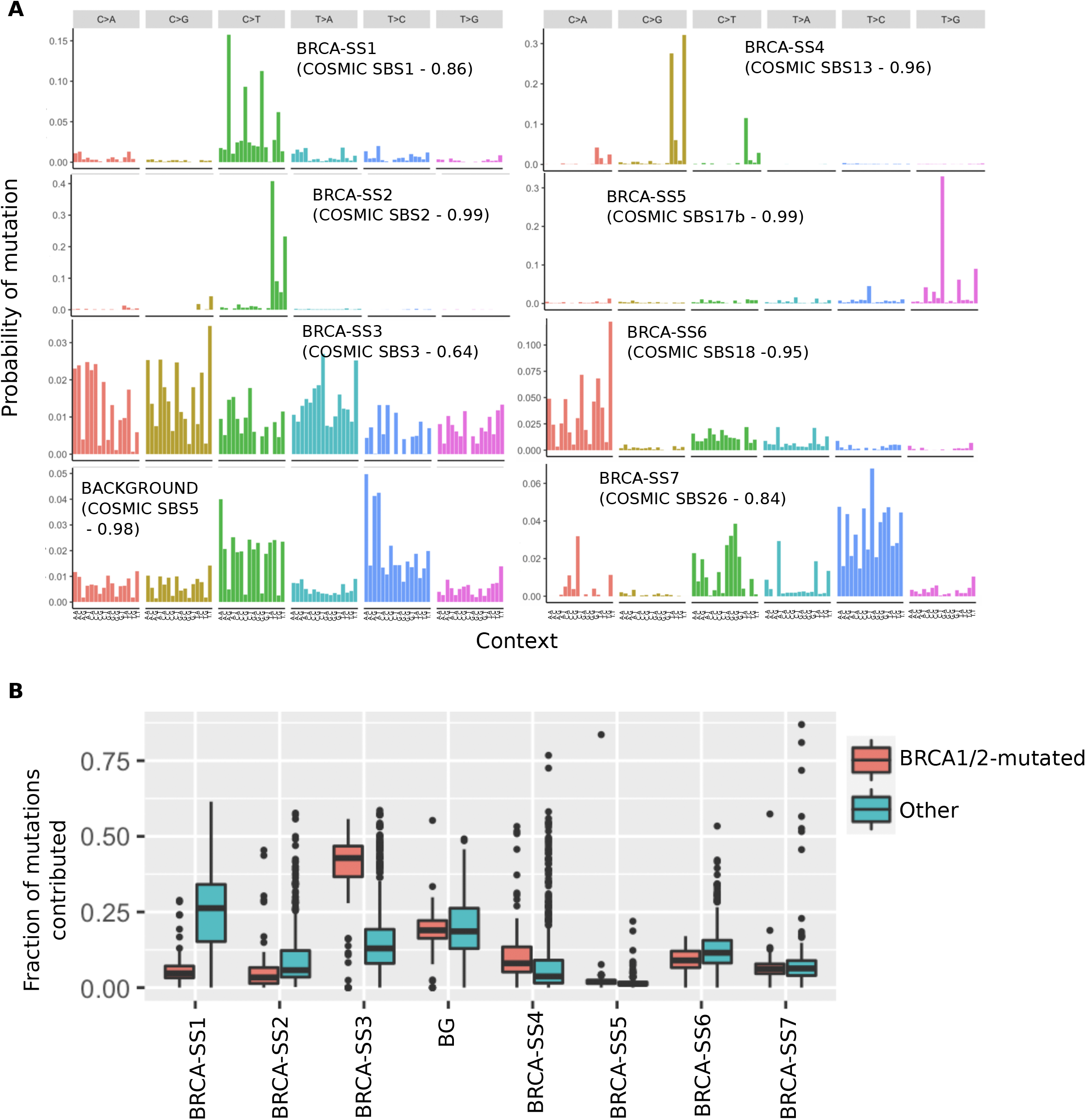
A) The 8 mutational signatures obtained by applying SparseSignatures to a dataset of 560 breast tumors. We report the number and correlation of the most similar (correlation higher than 0.70) corresponding signature from COSMIC (https://cancer.sanger.ac.uk/cosmic/signatures). Source data are provided in Supplementary Table 23. B) Boxplots showing fitted values for exposure to each of the 8 signatures obtained by SparseSignatures for BRCA1/2-mutated and non-mutated tumors. Boxplots represent the fraction of mutations per tumor (on the y-axis) contributed by the given signature (on the x-axis). Source data are provided in Supplementary Table 24.

We also examined the ability of SparseSignatures to accurately reconstruct signatures that occur in only a fraction of patients in the population. The simulated datasets for Simulation 3 were generated such that, in each dataset, some signatures were present in only a subset of patients. We found that SparseSignatures was able to reconstruct both rare and abundant signatures in these simulated datasets with high accuracy (Supplementary Figure 9A, Supplementary Table 10). In fact, SparseSignatures recovered rare signatures that were present in <35% of simulated patients with a median cosine similarity of 0.972; this is higher than the cosine similarities obtained by other methods (SigProfiler=0.881, signerR =0.913, SignatureAnalyzer=0.971) although these differences are not statistically significant. Similarly, SparseSignatures was able to accurately reconstruct signatures that contributed relatively few (<6,000) mutations to the dataset (Supplementary Figure 9B, Supplementary Table 10).

### SparseSignatures discovers well-differentiated signatures in pancreatic cancer data

We applied SparseSignatures to a dataset of patients affected by pancreatic cancer from PCAWG, including 147 curated whole genomes (Supplementary Table 11). Our goal was to discover mutational signatures that can be reconstructed with high accuracy and confidence. We therefore limited our analysis only to high-quality genomes with at least 1000 point mutations.

SparseSignatures discovered 8 signatures in addition to the background (Figure 3A, Supplementary Tables 12-13) along with their exposure values for each patient (Supplementary Table 14). We named these discovered signatures in the format “PC-SS”, for “Pancreatic Cancer - SparseSignatures”. We compared these signatures to literature on known mutational mechanisms and to the signatures described in the COSMIC database. Remarkably, most of the signatures can be associated with a known mutational process (Table 1). For example, PC-SS1 is caused by deamination of methylated cytosine in CpG contexts, and PC-SS2 and PC-SS4 by APOBEC enzymes.

We ran SigProfiler, SignatureAnalyzer, and signeR on the same dataset for comparison. All of these methods discovered 8 signatures (Supplementary Figures 10-12, Supplementary Tables 15-17). Compared to all other methods, SparseSignatures provides the best fit to the input data, in terms of overall residual error (Table 2) and also at the level of individual patients (Table 2, Supplementary Table 18), including patients with low as well as high mutation counts (Supplementary Figure 13). Further, the signatures discovered by SparseSignatures are sparser, and show the lowest similarity between signatures, indicating that they are more clearly differentiated from each other (Table 2). They also show the lowest similarity between the background and the non-background signatures, suggesting that the other sets contain noise due to imperfect separation of the background signature (Table 2).

This is supported by visual inspection of the signatures predicted by the four methods. The signatures predicted by SigProfiler and signeR appear to contain visible background noise (Supplementary Figures 10, 12). In addition, SPR7 (SigProfiler; Supplementary Figure 10) and SIP7 (signeR; Supplementary Figure 12) seem to result from imperfect separation of one of the well-known APOBEC mutagenesis signatures (PC-SS4), while SPR4 (SigProfiler; Supplementary Figure 10) and SIP5 (signeR; Supplementary Figure 12) show a low level of contamination with the CpG deamination signature (PC-SS1). The signatures produced by SignatureAnalyzer appear to lack the low level of background noise throughout, but show similar imperfect separation of APOBEC signatures in SIA3 and SIA7 (Supplementary Figure 11).

### Exposures predicted by SparseSignatures identify pancreatic cancer subtypes and correlate with clinical features

We next examined the exposure values produced by SparseSignatures for the background and 8 newly predicted signatures in pancreatic cancer samples. PC-SS1 (cytosine deamination at sites of CpG methylation) is the dominant signature, followed by the background signature and PC-SS6 (possibly reactive oxygen species) (Figure 3B, Supplementary Table 14).

We clustered all 147 tumors using CIMLR [32] based on these exposure values in order to identify subpopulations of tumors with similar mutagenic mechanisms. Using a bootstrap-based approach (Supplementary Methods) [33,34], we identified 10 clusters (Supplementary Figure 14, Supplementary Table 19) with different underlying exposures to the signatures (Figure 3D). C10 is high for PC-SS3 (likely representing defective homologous recombination-based DNA damage repair). The background signature is high in cluster 9, PC-SS6 is high in cluster 7, while cluster 8 seems to have high exposure to APOBEC signatures (PC-SS2 + PC-SS4).

The exposure-based clusters correlate with clinical features; cluster C1 with high PC-SS1 is enriched for females (Hypergeometric test p=0.0066), while cluster C10 has younger patients than the rest of the population (Wilcoxon test p=0.0333). Finally, patient relapse-free survival is significantly different between clusters, showing the potential clinical value of accurate signature discovery (Figure 3C). In contrast, the exposure values predicted by SignatureAnalyzer and signeR do not cluster patients into survival-associated subtypes, whereas clusters based on SigProfiler present a less significant association with survival (Supplementary Figure 15, Supplementary Table 20).

### SparseSignatures discovers a signature that characterizes BRCA-positive breast cancers

Finally, we applied SparseSignatures to a dataset of 560 breast tumors (ICGC Project BRCA-EU available from ICGC Data Portal https://icgc.org) (Supplementary Table 21). This dataset includes several different subtypes of breast cancer (118 triple-negative tumors, 293 ER+/HER2-tumors and 71 HER2+ tumors, as well as 36 tumors with BRCA1 and 39 tumors with BRCA2 mutations). Thus, this example illustrates the performance of SparseSignatures on a large and diverse dataset.

SparseSignatures discovers 7 well-differentiated signatures in addition to the background, all of which can be associated with known mutagenic processes in breast cancer (Figure 4A, Table 1, Supplementary Tables 22-23). As with pancreatic cancer, these results also include well-characterized mutational signatures associated with C>T deamination at CpG methylation sites and APOBEC enzymes. Moreover, SparseSignatures discovered a dense signature (BRCA-SS3), similar to COSMIC SBS3, that was significantly elevated in BRCA1/2 mutated tumors (Figure 4B, Supplementary Table 24, one-sided Wilcoxon test p < 2.2 × 10^−16^); this demonstrates its ability to recover signatures present in a small subset of tumors, as well as to recover dense signatures in addition to the background.

We compared the performance of SparseSignatures on this dataset to that of the other three methods. SigProfiler discovers 12 signatures in total (Supplementary Figure 16, Supplementary Table 25), allowing it to fit the dataset with a lower MSE than SparseSignatures; however, SparseSignatures still fits the counts of individual patients better (Table 2), regardless of the number of mutations in the tumor (Supplementary Figure 17, Supplementary Table 26); it also provides sparser, better differentiated signatures (Table 2). Moreover, while SparseSignatures, SignatureAnalyzer and signeR all discovered dense, BRCA-specific signatures close to SBS3 (see SIA3 and SIR3 in Supplementary Figures 18 and 19), sigProfiler did not.

SignatureAnalyzer also discovers 12 signatures (Supplementary Figure 18, Supplementary Table 27), but fits the data poorly (Table 2); moreover, it discovers a highly sparse signature that is not associated with any known mutagenic mechanism nor with any signature in COSMIC (SIA11, Supplementary Figure 18), and is likely to be an artifact owing to the tendency of this method to discover too many signatures. Finally, signeR discovers 7 signatures (Supplementary Figure 19, Supplementary Table 28) which are similar to those found by SparseSignatures. However, SparseSignatures fits the observed data better and presents sparser and better-differentiated signatures. Moreover, while SigProfiler and SignatureAnalyzer discover signatures similar to the background (SPR3 and SIA4 respectively in Supplementary Figures 17 and 18), signeR is unable to differentiate the background from the other signatures. Instead, it produces the signature SIR1, which is a mix of the CpG methylation signature and the background signature (Supplementary Figure 19; cosine similarity 0.86 with COSMIC Signature 1 and 0.72 with COSMIC Signature 5). This further validates our choice to fix the background signature in our method.

## Discussion

SparseSignatures is a novel approach designed to discover the best number of clearly differentiated mutational signatures with minimal background noise, which have robust statistical support by repeated cross-validation on unseen data points and are not likely due to overfitting.

Complementing its methodological innovations, SparseSignatures offers users the option to model a constant background signature. While users can supply a signature of their choice, we offer a background based on the COSMIC SBS5 signature, which, owing to its ubiquity in cancer and non-cancer tissues and cell lines, its correlation with age of diagnosis in cancers from multiple tissues [10], and its correlation with donor age in adult stem cells [22], has been hypothesized to represent clock-like mutational processes. Studies of human germline *de novo* mutations [23,24] and 1000 Genomes Project SNPs in different populations [23] show that the human germline mutational spectrum can be largely explained by SBS5 along with SBS1. We calculated a cosine similarity of 0.98 between SBS5 and the human germline mutational spectrum from trio studies [24], reinforcing the hypothesis that this signature represents a common spectrum of replication errors occuring in the normal course of cell division. Although the exact molecular causes of this signature are unknown, it may be a combination of several processes including proofreading errors by DNA polymerases and transcription-coupled repair [35].

While the use of the fixed background signature contributes to the strong performance of SparseSignatures in simulations (Figure 2), it must be treated with caution when applied to real data. Since the studies supporting this signature do not capture the full diversity of the human population, it may not represent understudied populations with equal accuracy. We anticipate that additional data from diverse human populations will help improve our model. There may also exist some patients where this signature is altered, due to variation in its underlying processes, e.g. proofreading enzymes. Along with manual examination of discovered signatures, careful examination of residual error and per-sample correlation metrics can diagnose whether some samples in a dataset are not fitted well due to the assumptions of the model. In our analyses on pancreatic and breast tumors, the background signature was consistently abundant (Supplementary Table 14, Supplementary Table 24, Figure 3B, Figure 4B), and SparseSignatures presented extremely accurate reconstruction of all individual samples (Supplementary Table 18, Supplementary Table 26, Table 2), as well as low mean squared error across the entire datasets (Table 2).

The density and complexity of the SBS5/background signature renders it particularly difficult to distinguish *de novo* from the low number of samples typically available in cancer studies. If not distinguished accurately, components of this signature can be mixed with other signatures, leading to inaccurate results. An example can be observed in our analysis of breast cancer, where signeR was unable to distinguish the background signature, instead combining it with the well-known SBS1 (Supplementary Figure 19). Although caution is necessary, we believe that fixing the background signature is a useful option that can benefit many studies.

Further, our method supports the discovery of sparse signatures by applying a LASSO penalty to the signatures matrix. We also offer the option to apply a similar penalty to regularize the exposure matrix, since it is also reasonable to believe that only a limited number of mutational processes will be active in each patient. However, this option presents a high computational cost, and our experiments thus far show that it produces a relatively minor improvement in results. We are currently incorporating an option to allow multiple fixed signatures in addition to the background, such as the signature of cytosine deamination or other signatures that are known to exist in the cancer type being studied, as suggested by previous literature [26]. Future work could also be directed at incorporating indels and doublet base substitutions [11], especially when larger datasets become available to support analyses of these rarer events.

Multiple experiments on simulated data show that SparseSignatures outperforms current state-of-the-art methods. It provides the most accurate and least ambiguous estimation of the number of signatures, and reconstructs the original signatures and exposures most accurately. In comparison, other methods tend to discover too many signatures or retain noise in the discovered signatures. Further, we have applied SparseSignatures to whole genome sequences from 147 pancreatic tumors and 560 breast tumors. Compared to other methods, we successfully obtain a good fit to the observed data, while at the same time obtaining signatures that are sparse, differentiated, have reduced noise, and are attributable to known biological processes while at the same time preventing overfitting. The signatures discovered by SparseSignatures are predictive of patient survival in pancreatic cancer (Figure 3C), and associated with known biological subtypes in breast cancer (Figure 4B). We anticipate that the availability of larger datasets comprising curated, uniformly processed whole genome sequences may allow us to validate those signatures and discover new ones.

In conclusion, the small number of highly specific, differentiated signatures discovered by SparseSignatures leads us to predict that whole genome sequencing of individual cancers and their classification on the basis of signatures, including the background, may become much more easily interpretable and possibly useful in a clinical context. For example, strong contribution of CpG methylation versus background in a patient suggests that methylation changes have been more important for the growth of the cancer and that overall cellular turnover (associated with background) may have been modest, suggesting that DNA replication inhibitors may be less effective than gene regulatory therapy for such patients. We suggest that future work be directed at greater numbers of patients for whole genome sequencing and the simultaneous collection of other omic data to connect mutagenesis with molecular phenotype and eventually mechanistic cause.

## Methods

### Mathematical Framework for Mutational Signature Discovery

The mathematical framework developed for signature extraction [4] is as follows. First, all point mutations are classified into 6 groups (C>A, C>G, C>T, T>A, T>C, T>G; the original pyrimidine base is listed first). Then, these are subdivided into 16 × 6 = 96 categories based on the 16 possible combinations of 5’ and 3’ flanking bases. Each tumor sample is described by the count of mutations in each of the 96 categories. This forms a count matrix M, where the rows are the tumor samples and the columns are the 96 categories.

Signature extraction aims to decompose M into the multiplication of two low-rank matrices: the exposure matrix α and the signature matrix β.

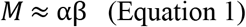

Here, α is the exposure matrix with one row per tumor and K columns, and β is the signature matrix with K rows and 96 columns. K is the number of signatures. Each row of β represents a signature, and each row of α represents the exposure of a single tumor to all K signatures, i.e. the number of mutations contributed by each signature to that tumor. In NMF, this equation is solved for α and β by minimizing the squared residual error (some methods use Kullback–Leibler divergence instead) while constraining all elements of α and β to be non-negative.

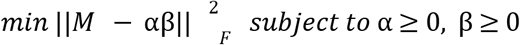

### Improvements to the NMF framework in SparseSignatures

In SparseSignatures, we incorporate a background signature by modifying Equation (1) as follows:

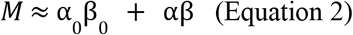

Here, β_0_ is the known ‘background’ signature of point mutations caused by replication errors during cell division, and α_0_ is the vector of exposures of all tumors to that signature. The dimensions of α_0_ are (number of tumors x 1) and the dimensions of β_0_ are 1 × 96.

To enforce sparsity in the discovered signatures, we use the LASSO [14]. This is done by adding an additional regularization term to the cost function to be minimized:

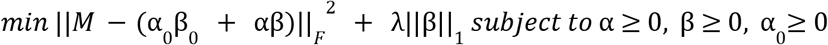

The parameter λ controls the extent to which sparsity is encouraged in the signature matrix β. If the value of λ is set too low, it is ineffective, whereas if it is set too high, the signatures are forced to be too sparse and no longer accurately fit the data.

It should be noted that unlike the standard LASSO, the objective function we minimize here is non-convex. But it is bi-convex (convex in α with β fixed and vice-versa). Hence the alternating algorithm described below is natural and yields good solutions. A standard issue with all NMF algorithms is non-identifiability: if we scale βby *c* andα by 1/*c*, the product αβ remains unchanged. One can change the relative magnitudes of α and β at convergence by changing their relative magnitudes at initialization. To remove this ambiguity, we initialize β so that each row (signature) sums to 1. The choice of 1 is not important: if we had instead initialized β so that each row sums to *c*, the signatures we obtain at algorithm convergence would be equivalent (up to proportionality) to those obtained by initializing β with all rows summing to 1 and λ set to λ/*c*.

### Implementation of SparseSignatures

SparseSignatures discovers mutational signatures by following the steps below.

**Step 1:** Build the Count Matrix M by counting the number of mutations of each of the 96 categories in each sample.

**Step 2**: Remove samples with less than a minimum number of mutations. In the analysis described in this paper, we have used a minimum number of 1000 mutations per tumor genome.

**Step 3:** Choose a range of values to test for K (number of signatures) and λ (level of sparsity).

**Step 4:** For each value of K in the chosen range, obtain a set of K initial signatures using repeated NMF

[36] to obtain a more robust estimation. This is an initial value for the matrix β. We use these NMF results as a starting point (although other starting points such as randomly generated signatures may also be chosen) and further refine the signatures. In practice, the final discovered signatures are often very different from those produced by the initial NMF.

**Step 5:** For each pair of parameter values (K and λ), perform cross-validation as follows [27]:

**5a**. Randomly select a given percentage of cells from M. Based on simulations (Supplementary Methods, Supplementary Table 29), we currently use 1% of the points in the dataset for cross-validation; however, the method appears robust to large variations in this value.

**5b**. Replace the values in those cells with 0.

**5c**. Consider the NMF results for the chosen value of K as an initial value of β. Add the background signature (β_0_). Then use an iterative approach to discover signatures with sparsity. Each iteration involves two steps:

**5c(i)**. While keeping fixed the values of β_0_ and β, fit α_0_ and α by minimizing:

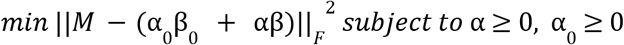

**5c(ii)**. While keeping fixed the values of β_0_, α_0_ and α, fit β by minimizing:

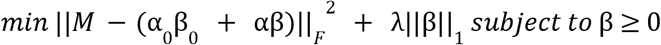

These steps are repeated for a number of iterations (set to 20 by default; in all our experiments we found that this was sufficient to reach convergence).

**5d**. Use the obtained signatures to predict the values for the cells that were set to 0 (we do this by calculating the matrix α_0_ β_0_ + αβ and taking the entries corresponding to the cross-validation cells).

Then replace the values in these cells with the predicted values and repeat step 5c. We repeat step 5c a number of times (set to 5 by default), each time discovering signatures and then replacing the values of the cross-validation cells by the predicted values. After each iteration, the predictions improve, as the algorithm converges, making the mean squared errors used in the next step more stable.

**5e**. At the last iteration of step 5d, measure the mean squared error (MSE) of the prediction.

**5f**. Repeat the entire cross-validation procedure (steps 5a-5d) a number of times (set to 10 by default) and calculate the MSE for all cross-validations. Since we randomly select a different set of cells for cross-validation each time, this allows us to obtain a robust measure of MSE.

**Step 6:** Choose the values of K and λ that correspond to the lowest MSE in most of the cross-validations. **Step 7:** Using the selected values for K and λ, repeat sparse signature discovery (step 5c) on the complete matrix M (without replacing any cells with 0). This generates the final values of α_0_, α and β.

### Background signature

SparseSignatures offers two preset options for the background signature. The first is derived from the germline mutation spectrum calculated by [24]. To validate this, we independently calculated the germline mutational spectrum using whole-genome sequencing data from normal tissue samples (see Supplementary Methods for details), and the spectrum thus obtained had a high cosine similarity of 0.98 with that calculated by [24]. We then adjusted the rates of ACG>ATG, CCG>CTG, GCG>GTG and TCG>TTG mutations to be equal to the rates of ACA>ATA, CCA>CTA, GCA>GTA and TCA>TTA mutations respectively, in order to separate the effects of DNA methylation from the background signature. The second is derived from the SBS5 signature in COSMIC v3, which has been found across diverse human tumor types and has been associated with cellular turnover and aging. Here, we once again empirically adjusted the rates of ACG>ATG, CCG>CTG, GCG>GTG and TCG>TTG mutations. For the experiments described here, we used the germline-derived signature.

### Definition of the λ parameter

This parameter tunes the desired level of regularization to be obtained by LASSO. For any analysis by LASSO, one can compute a maximal value of the LASSO penalty after which all the coefficients of the regression get shrunk to zero [37]. As this maximal value can vary depending on the problem, our λ parameter represents the fraction of the actual maximal value to be used. Values closer to 1 result in higher regularization.

### Simulations

We performed 6 simulated experiments all including 50 simulated datasets. The first five simulations included 4 signatures and simulation 6 included 8 signatures. In Simulation 1, we used real data to perform simulations and specifically we considered 116 curated WGS data of prostate cancer samples obtained from PCAWG (https://dcc.icgc.org/pcawg) with at least 1000 mutations, and selected a set of 4 signatures from COSMIC known to be active in prostate [11]; we then used deconstructSigs [38] to fit such signatures on the data and generate their assignments to samples. Furthermore, we performed three additional experiments (Simulations 2-4) where we randomly selected signatures from COSMIC, considering all of them (Simulation 2) as well as the subset of dense (Simulation 3) or sparse (Simulation 4) signatures; we then generated random assignments of such signatures to samples, for a total of 100 samples per experiment. In Simulation 5, we used the same settings of Simulation 1, but including both additive and subtractive noise; finally in Simulation 6, we used a similar configuration to the one of Simulation 2, this time including a total of 8 signatures chosen randomly from COSMIC database.

We ran four methods for *de novo* signature discovery (SparseSignatures, SigProfiler, SignatureAnalyzer, and signeR) on each of the 50 datasets and evaluated their performance. These methods were executed with the configurations suggested by the authors in the respective manuscripts. Specifically, SignatureAnalyzer was performed 10 times and the solution with best posterior was chosen; SigProfiler pipeline was performed 10 times with 100 iterations each. Details are provided in the Supplementary Methods. To evaluate the accuracy with which discovered signatures reconstructed the original signatures, we matched each input signature to its closest discovered signature and evaluated the match by mean squared error. We then also measured the mean squared error between the exposure values of the input signature and the discovered exposure values for its most similar discovered signature. Further details are given in the Supplementary Methods.

### Real data

We obtained a dataset of point mutations in pancreatic tumors from ICGC (see Supplementary Table 11 for the full list of samples). We selected only whole-genome sequencing data and removed samples with less than 1000 point mutations. After this preprocessing, a total of 147 samples remained. Further, we obtained a dataset of point mutations from ICGC (see Supplementary Table 21 for the full list of samples) comprising whole-genome sequencing for a total of 560 samples.

### Software

The experiments carried out in this paper were performed using the SparseSignatures v2.0.0 R package and R version 4.0.3. The software is available for download on Bioconductor at https://bioconductor.org/packages/release/bioc/html/SparseSignatures.html. This package in its current version makes use of external R packages NMF v0.21.0 [39], nnls v1.4 and nnlasso v0.3.

## Data Access

The whole-genome sequencing data used in this study is publicly available and was downloaded from https://dcc.icgc.org/search.

## Acknowledgments

This work was supported by an R01 grant to A.S. (NIH/NCI) and gift funding from the BRCA Foundation. A.L. was supported by a Young Investigator Award from the BRCA Foundation. D.R. was partially supported by a Bicocca 2020 Starting Grant and by a Premio Giovani Talenti dell’Università degli Studi di Milano-Bicocca. The results published here are based in part upon data generated by the Pan-Cancer Analysis of Whole Genomes (PCAWG) Research Network (https://dcc.icgc.org/pcawg).

## Supplementary Material Legends

**Supplementary Methods**. Supplementary information of the experiments presented in the manuscript.

**Supplementary Figure 1**. A) Average mutational counts for 116 simulated patients in each of 96 mutational categories. This dataset is one of 50 datasets simulated as part of Simulation 1. Error bars represent standard deviation. B) 4 original signatures in the simulated dataset. C) 4 signatures deciphered by SparseSignatures from the simulated dataset. D) 4 signatures deciphered by SparseSignatures from the simulated dataset, without the fixed background. Source data are provided in Supplementary Table 3.

**Supplementary Figure 2**. A) 4 signatures deciphered by SigProfiler from the simulated dataset shown in Supplementary Figure 1A. B) 4 signatures deciphered by SignatureAnalyzer from the simulated dataset. C) 4 signatures deciphered by signeR from the simulated dataset. Source data are provided in Supplementary Table 3.

**Supplementary Figure 3**. Comparison between SparseSignatures and other methods on simulated data when the correct number of signatures is known. A) Box plots showing the residual error for the solutions produced by each method, over 50 simulations. Residual error was measured as the mean squared error (MSE) in reconstructing the original count matrix. B) Box plots showing the fraction of variance in the count matrix explained by the solutions produced by each method, over 50 simulations. (C) Box plots showing the cosine similarity of reconstructing the 3 non-background input signatures, over 50 simulations. D) Box plots showing the mean squared error in reconstructing the exposure values for the 3 non-background input signatures, over 50 simulations. E) Box plots showing the sparsity of the signatures produced by each method, over 50 simulations. Sparsity was measured as the fraction of cells in the signature matrix whose value is <10^−3^. SS: SparseSignatures. SP: SigProfiler. SA: SignatureAnalyzer. SR: signeR. Source data are provided in Supplementary Table 4.

**Supplementary Figure 4**. Comparison between SparseSignatures and other methods on simulated data with both additive and subtractive noise (see Supplementary Methods for details). A) Bar and line plot showing, for each method, the number of simulation runs in which it selected each value of K (number of signatures). The x-axis shows values of K and the y-axis shows the number of times each value was selected. Each method was run on 50 simulated datasets. In all cases, the correct value of K was 4. B) Box plots showing the residual error for the solutions produced by each method, over 50 simulations. Residual error was measured as the mean squared error (MSE) in reconstructing the original count matrix. C) Box plots showing the fraction of variance in the count matrix explained by the solutions produced by each method, over 50 simulations. D) Box plots showing the cosine similarity of reconstructing the 3 non-background input signatures, over 50 simulations. E) Box plots showing the mean squared error in reconstructing the exposure values for the 3 non-background input signatures, over 50 simulations. F) Box plots showing the sparsity of the signatures produced by each method, over 50 simulations. Sparsity was measured as the fraction of cells in the signature matrix whose value is <10^−3^. SS: SparseSignatures. SP: SigProfiler. SA: SignatureAnalyzer. SR: signeR. Source data are provided in Supplementary Table 5.

**Supplementary Figure 5**. Comparison between SparseSignatures and other methods on simulated data generated from 4 randomly selected COSMIC signatures. A) Bar and line plot showing, for each method, the number of simulation runs in which it selected each value of K (number of signatures). The x-axis shows values of K and the y-axis shows the number of times each value was selected. Each method was run on 50 simulated datasets. In all cases, the correct value of K was 4. B) Box plots showing the residual error for the solutions produced by each method, over 50 simulations. Residual error was measured as the mean squared error (MSE) in reconstructing the original count matrix. C) Box plots showing the fraction of variance in the count matrix explained by the solutions produced by each method, over 50 simulations. D) Box plots showing the cosine similarity of reconstructing the 3 non-background input signatures, over 50 simulations. E) Box plots showing the mean squared error in reconstructing the exposure values for the 3 non-background input signatures, over 50 simulations. F) Box plots showing the sparsity of the signatures produced by each method, over 50 simulations. Sparsity was measured as the fraction of cells in the signature matrix whose value is <10^−3^. SS: SparseSignatures. SP: SigProfiler. SA: SignatureAnalyzer. SR: signeR. Source data are provided in Supplementary Table 6.

**Supplementary Figure 6**. Comparison between SparseSignatures and other methods on simulated data generated from 4 randomly selected dense COSMIC signatures. A) Bar and line plot showing, for each method, the number of simulations in which it selected each value of K (number of signatures). The x-axis shows values of K and the y-axis shows the number of times each value was selected. Each method was run on 50 simulated datasets. In all cases, the correct value of K was 4. B) Box plots showing the residual error for the solutions produced by each method, over 50 simulations. Residual error was measured as the mean squared error (MSE) in reconstructing the original count matrix. C) Box plots showing the fraction of variance in the count matrix explained by the solutions produced by each method, over 50 simulations. D) Box plots showing the mean squared error in reconstructing the 3 non-background input signatures, over 50 simulations. E) Box plots showing the mean squared error in reconstructing the exposure values for the 3 non-background input signatures, over 50 simulations. F) Box plots showing the sparsity of the signatures produced by each method, over 50 simulations. Sparsity was measured as the fraction of cells in the signature matrix whose value is <10^−3^. SS: SparseSignatures. SP: SigProfiler. SA: SignatureAnalyzer. SR: signeR. Source data are provided in Supplementary Table 7.

**Supplementary Figure 7**. Comparison between SparseSignatures and other methods on simulated data generated from 4 randomly selected sparse COSMIC signatures. A) Bar and line plot showing, for each method, the number of simulations in which it selected each value of K (number of signatures). The x-axis shows values of K and the y-axis shows the number of times each value was selected. Each method was run on 50 simulated datasets. In all cases, the correct value of K was 4. B) Box plots showing the residual error for the solutions produced by each method, over 50 simulations. Residual error was measured as the mean squared error (MSE) in reconstructing the original count matrix. C) Box plots showing the fraction of variance in the count matrix explained by the solutions produced by each method, over 50 simulations. D) Box plots showing the cosine similarity of reconstructing the 3 non-background input signatures, over 50 simulations. E) Box plots showing the mean squared error in reconstructing the exposure values for the 3 non-background input signatures, over 50 simulations. F) Box plots showing the sparsity of the signatures produced by each method, over 50 simulations. Sparsity was measured as the fraction of cells in the signature matrix whose value is <10^−3^. SS: SparseSignatures. SP: SigProfiler. SA: SignatureAnalyzer. SR: signeR. Source data are provided in Supplementary Table 8.

**Supplementary Figure 8**. Comparison between SparseSignatures and other methods on simulated data generated from 8 randomly selected COSMIC signatures. A) Bar and line plot showing, for each method, the number of simulations in which it selected each value of K (number of signatures). The x-axis shows values of K and the y-axis shows the number of times each value was selected. Each method was run on 50 simulated datasets. In all cases, the correct value of K was 8. B) Box plots showing the residual error for the solutions produced by each method, over 50 simulations. Residual error was measured as the mean squared error (MSE) in reconstructing the original count matrix. C) Box plots showing the fraction of variance in the count matrix explained by the solutions produced by each method, over 50 simulations. D) Box plots showing the cosine similarity of reconstructing the 7 non-background input signatures, over 50 simulations. E) Box plots showing the mean squared error in reconstructing the exposure values for the 7 non-background input signatures, over 50 simulations. F) Box plots showing the sparsity of the signatures produced by each method, over 50 simulations. Sparsity was measured as the fraction of cells in the signature matrix whose value is <10^−3^. SS: SparseSignatures. SP: SigProfiler. SA: SignatureAnalyzer. SR: signeR. Source data are provided in Supplementary Table 9.

**Supplementary Figure 9**. Performance of SparseSignatures and other methods at reconstructing rare signatures, using the same data as Supplementary Figure 5. A) Boxplots showing the cosine similarity of signature reconstruction for signatures, separated by the fraction of patients in the population in which the signature is present. B) Boxplots showing the cosine similarity of signature reconstruction for signatures, separated by the number of mutations contributed by the signature in the overall dataset. Source data are provided in Supplementary Table 10.

**Supplementary Figure 10**. 8 signatures predicted by SigProfiler on 147 pancreatic tumors. Source data are provided in Supplementary Table 15.

**Supplementary Figure 11**. 8 signatures predicted by SignatureAnalyzer on 147 pancreatic tumors. Source data are provided in Supplementary Table 16.

**Supplementary Figure 12**. 8 signatures predicted by signeR on 147 pancreatic tumors. Source data are provided in Supplementary Table 17.

**Supplementary Figure 13**. Boxplots representing the Pearson Correlation between observed and predicted mutation counts in 96 categories, for individual patients in the dataset of 147 pancreatic tumors. The x-axis shows the total number of mutations in the tumor. SS: SparseSignatures. SP: SigProfiler. SA: SignatureAnalyzer. SR: signeR. Source data are provided in Supplementary Table 18.

**Supplementary Figure 14**. A) CIMLR was first run on the original dataset; then clustering was repeated 100 times on datasets generated by bootstrap resampling. The figure reports mean normalized mutual information (NMI) between cluster assignments across the bootstraps; higher values indicate stable results. B) CIMLR number of clusters for SparseSignatures.

**Supplementary Figure 15**. A) Survival curves for pancreatic cancer patients, divided into CIMLR clusters based on SigProfiler results. B) Survival curves for pancreatic cancer patients, divided into CIMLR clusters based on SignatureAnalyzer results. C) Survival curves for pancreatic cancer patients, divided into CIMLR clusters based on signeR results. Source data are provided in Supplementary Table 20.

**Supplementary Figure 16**. 12 signatures predicted by SigProfiler on 560 breast tumors. Source data are provided in Supplementary Table 25.

**Supplementary Figure 17**. Boxplots representing the Pearson Correlation between observed and predicted mutation counts in 96 categories, for individual patients in the dataset of 560 breast tumors. The x-axis shows the total number of mutations in the tumor. SS: SparseSignatures. SP: SigProfiler. SA: SignatureAnalyzer. SR: signeR. Source data are provided in Supplementary Table 26.

**Supplementary Figure 18**. 12 signatures predicted by SignatureAnalyzer on 560 breast tumors. Source data are provided in Supplementary Table 27.

**Supplementary Figure 19**. 7 signatures predicted by signeR on 560 breast tumors. Source data are provided in Supplementary Table 28.

**Supplementary Table 1**. Preset background signatures offered with SparseSignatures.

**Supplementary Table 2**. Performance metrics for signature discovery methods applied to simulated data (Simulation 1).

**Supplementary Table 3**. Average mutational counts, true signatures, and discovered signatures for one simulated dataset in Simulation 1.

**Supplementary Table 4**. Performance metrics for signature discovery methods applied to simulated data (Simulation 1) given the correct number of signatures.

**Supplementary Table 5**. Performance metrics for signature discovery methods applied to simulated data (Simulation 2).

**Supplementary Table 6**. Performance metrics for signature discovery methods applied to simulated data (Simulation 3).

**Supplementary Table 7**. Performance metrics for signature discovery methods applied to simulated data (Simulation 4).

**Supplementary Table 8**. Performance metrics for signature discovery methods applied to simulated data (Simulation 5).

**Supplementary Table 9**. Performance metrics for signature discovery methods applied to simulated data (Simulation 6).

**Supplementary Table 10**. Cosine similarity between non-background reconstructed signatures and original signatures in Simulation 3.

**Supplementary Table 11**. List of 147 Pancreatic cancer samples used for signature discovery.

**Supplementary Table 12**. Results of cross-validation to choose the best values of K and λ on pancreatic cancer data, using 1% of the cells in the matrix for cross-validation. We tested values of K ranging from 2 to 18 and values of lambda of 0.01, 0.025, 0.05, 0.075 and 0.1. Cross-validation was repeated 500 times with 5 restarts each. The entries in the table represent the median mean square error (MSE) in fitting the unseen data points across the 500 repetitions.

**Supplementary Table 13**. 9 signatures (including the background signature) discovered by applying SparseSignatures to pancreatic cancer data.

**Supplementary Table 14**. Fitted values for exposure to each of the 9 signatures (including the background signature) discovered by applying SparseSignatures to pancreatic cancer data, of each of the 147 whole genomes in the dataset.

**Supplementary Table 15**. 8 signatures discovered by applying SigProfiler to pancreatic cancer data.

**Supplementary Table 16**. 8 signatures discovered by applying SignatureAnalyzer to pancreatic cancer data.

**Supplementary Table 17**. 8 signatures discovered by applying signeR to pancreatic cancer data.

**Supplementary Table 18**. Mean correlation of observed and predicted counts for each of the 147 pancreatic cancer tumors.

**Supplementary Table 19**. Cluster assignments generated by CIMLR for each pancreatic tumor sample.

**Supplementary Table 20**. Number of patients at risk at each time point, according to various clusters defined by predicted exposures, in survival curves of 147 pancreatic cancer patients.

**Supplementary Table 21**. List of 560 Breast cancer samples used for signature discovery.

**Supplementary Table 22**. Results of cross-validation to choose the best values of K and λ on breast cancer data, using 1% of the cells in the matrix for cross-validation. We tested values of K ranging from 2 to 18 and values of lambda of 0.01, 0.025, 0.05, 0.075 and 0.1. Cross-validation was repeated 500 times with 5 restarts each. The entries in the table represent the median mean square error (MSE) in fitting the unseen data points across the 500 repetitions.

**Supplementary Table 23**. 8 signatures (including the background signature) discovered by applying SparseSignatures to breast cancer data.

**Supplementary Table 24**. Fitted values for exposure to each of the 8 signatures (including the background signature) discovered by applying SparseSignatures to breast cancer data, of each of the 560 whole genomes in the dataset.

**Supplementary Table 25**. 12 signatures discovered by applying SigProfiler to breast cancer data.

**Supplementary Table 26**. Pearson correlation between the true and reconstructed counts of individual breast tumor samples.

**Supplementary Table 27**. 12 signatures discovered by applying SignatureAnalyzer to breast cancer data.

**Supplementary Table 28**. 7 signatures discovered by applying signeR to breast cancer data.

**Supplementary Table 29**. Results of cross-validation to choose the best values of K on simulated data, using 0.1%, 1%, and 10% of the cells in the matrix M for cross-validation. Cross-validation was repeated 100 times for each percentage of cells. The entries in the table represent the number of times (over 100 repetitions) when a given value of K was chosen as optimal, based on it having the lowest median mean square error (MSE). The true value of K is 5.

