## Supplementary figures and images for "De Novo Mutational Signature Discovery in Tumor Genomes using SparseSignatures"

### SupplementaryFigure1.pdf

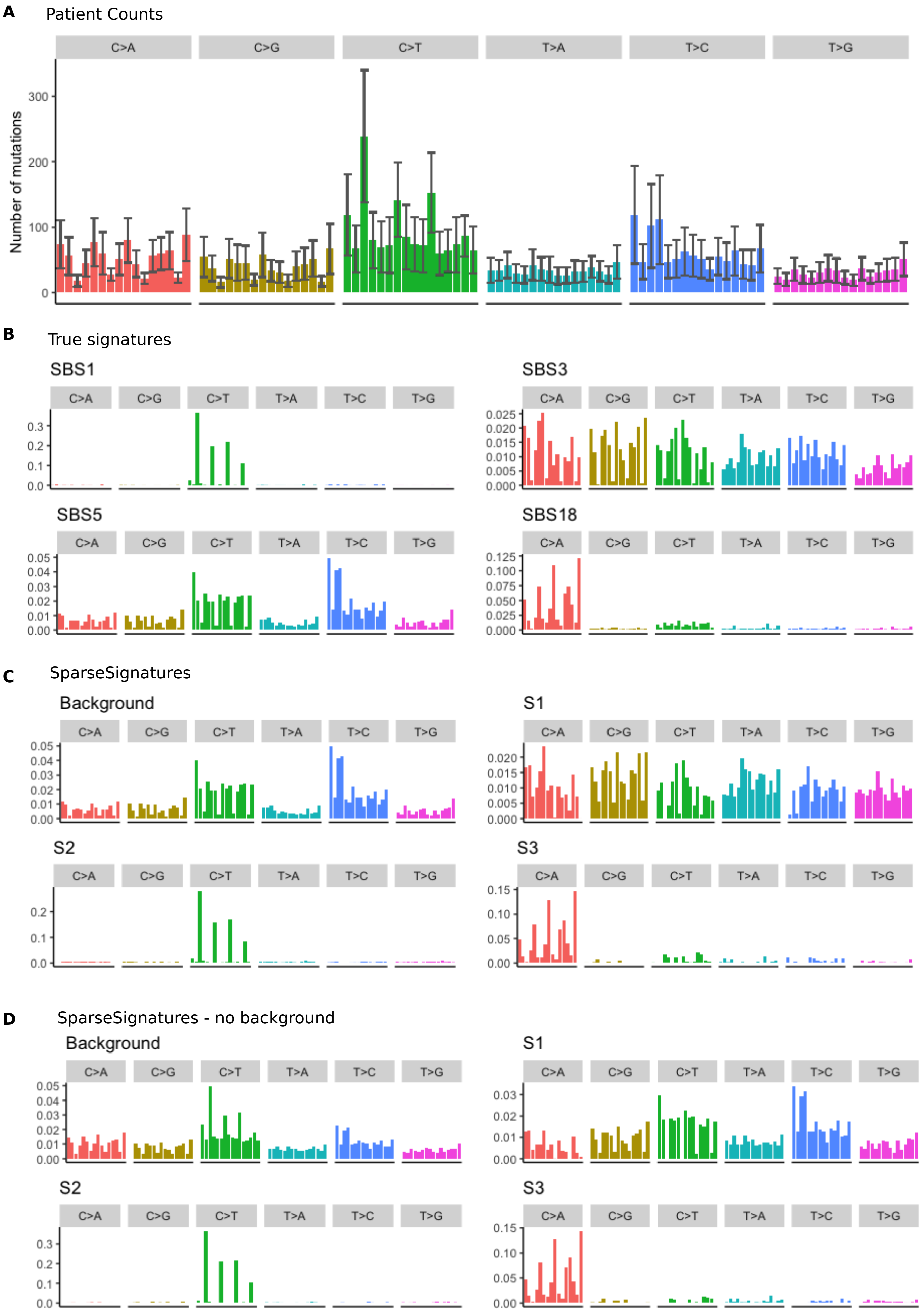

### SupplementaryFigure2.pdf

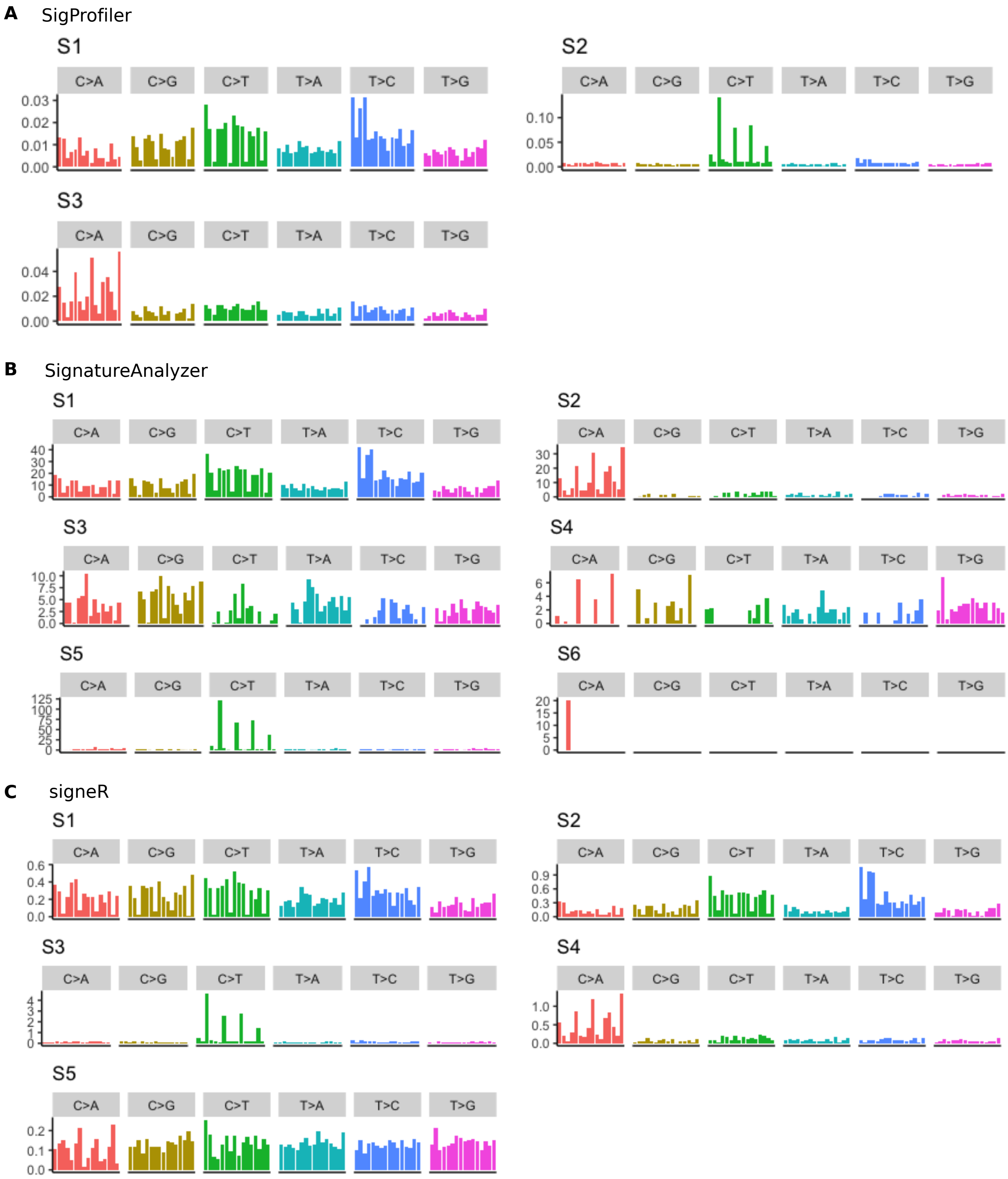

### SupplementaryFigure3.pdf

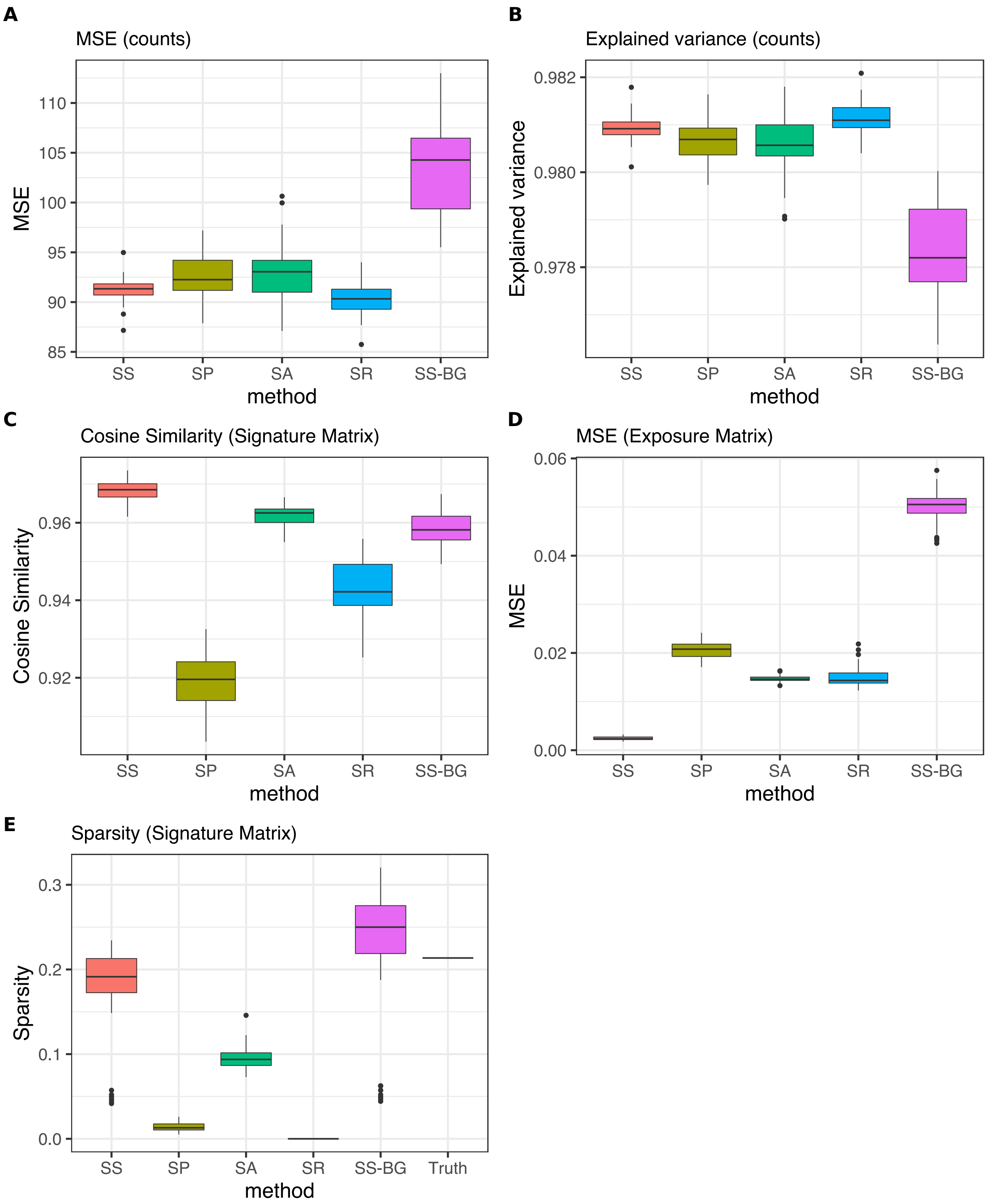

### SupplementaryFigure4.pdf

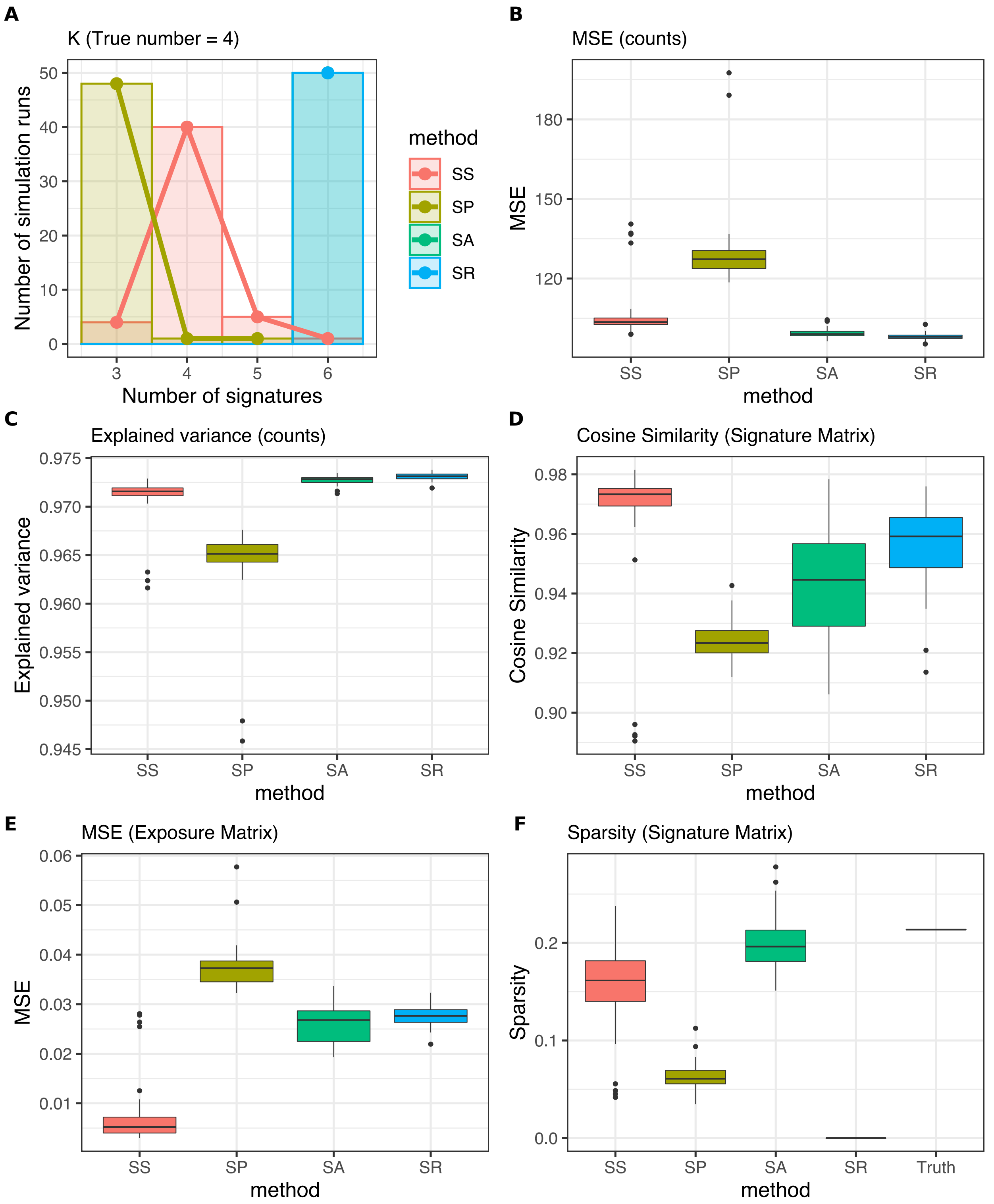

### SupplementaryFigure5.pdf

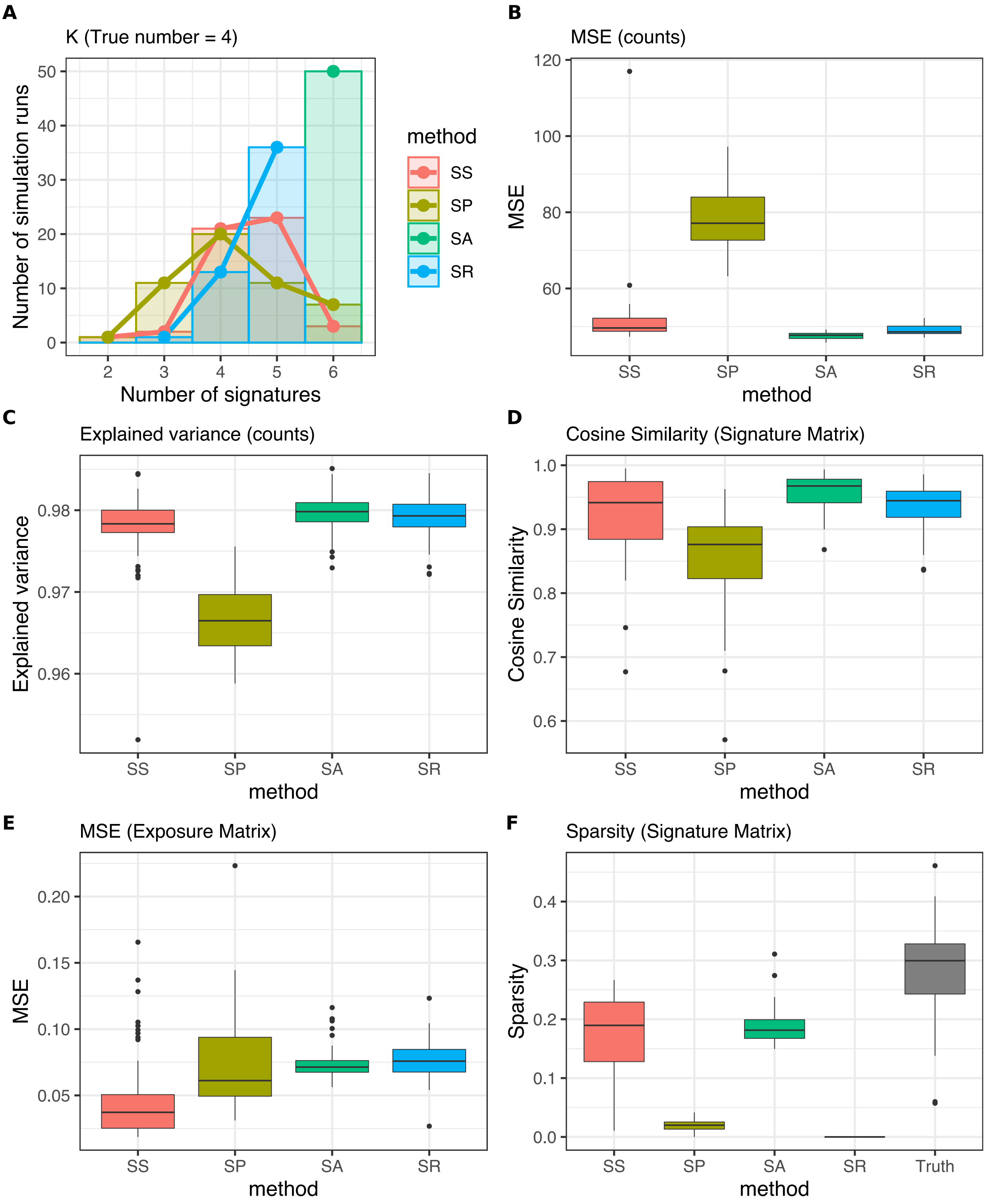

### SupplementaryFigure6.pdf

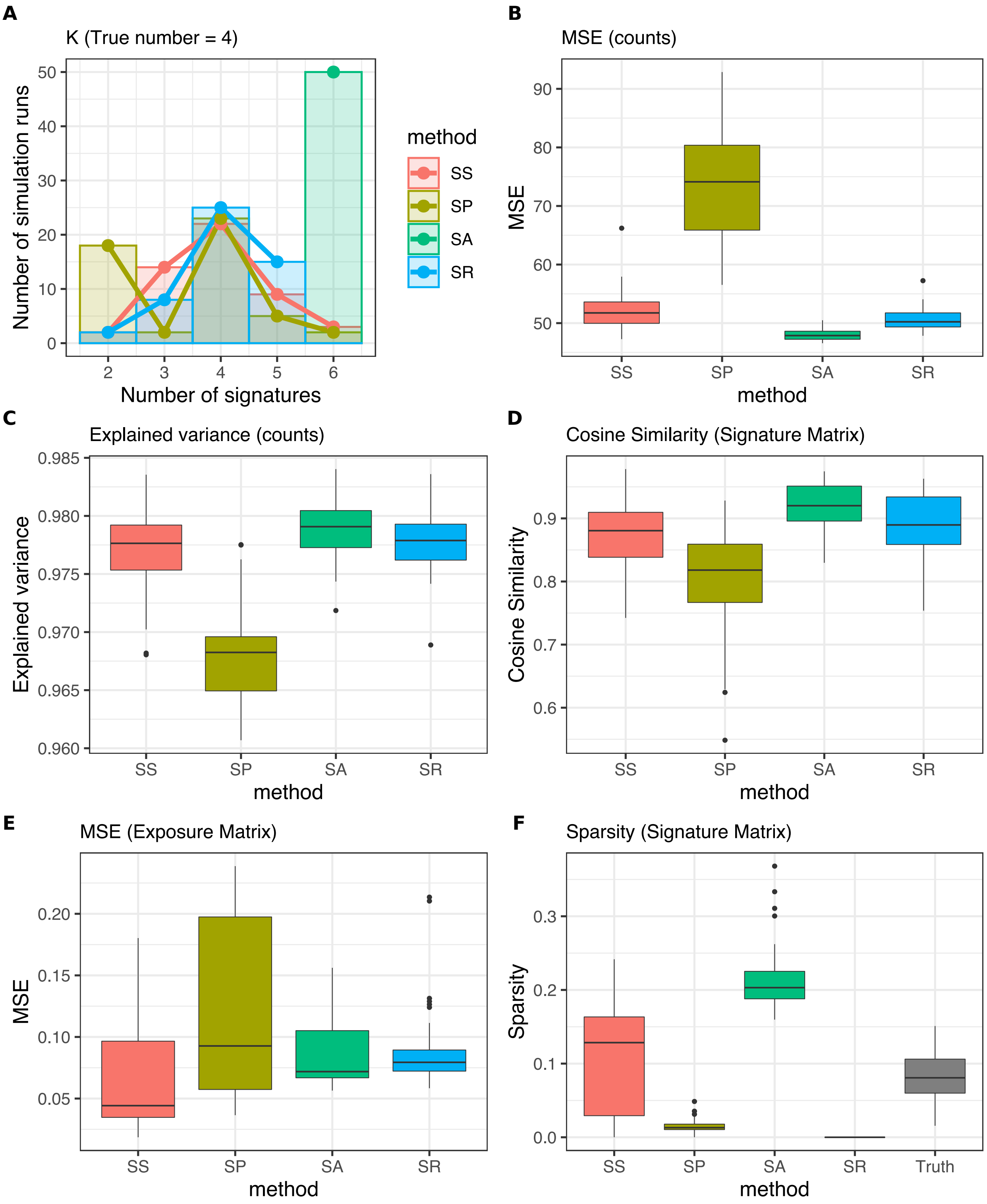

### SupplementaryFigure7.pdf

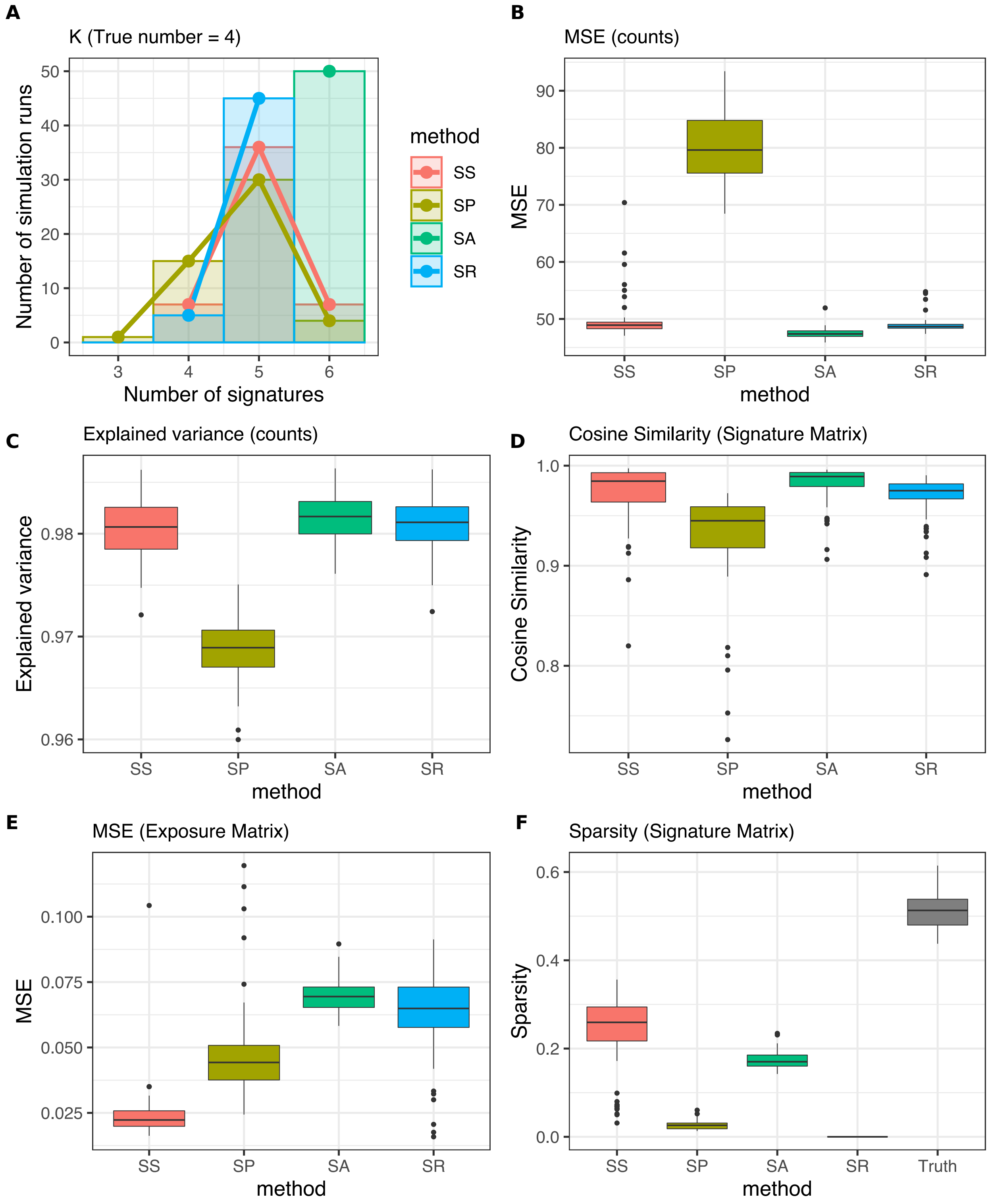

### SupplementaryFigure8.pdf

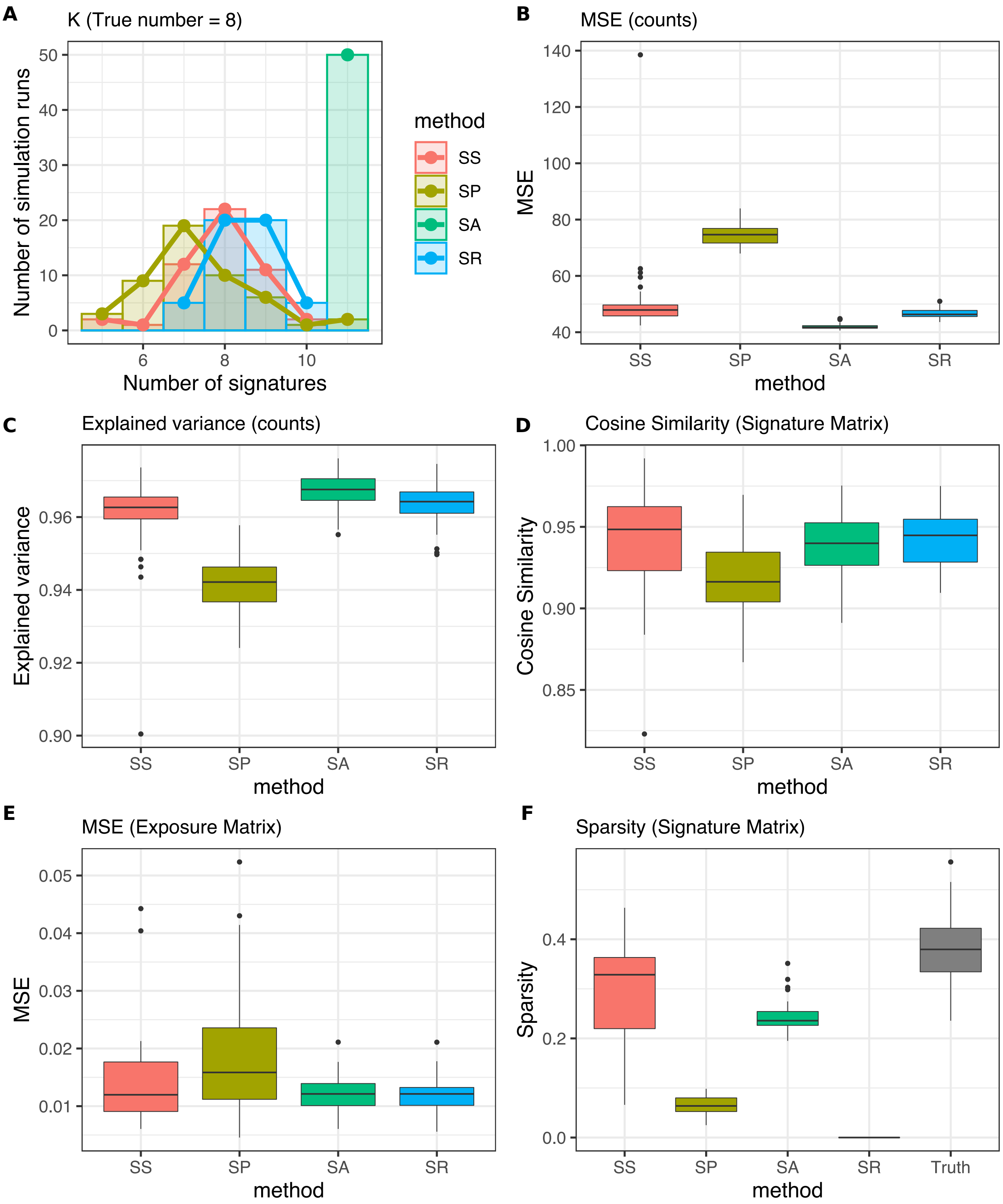

### SupplementaryFigure9.pdf

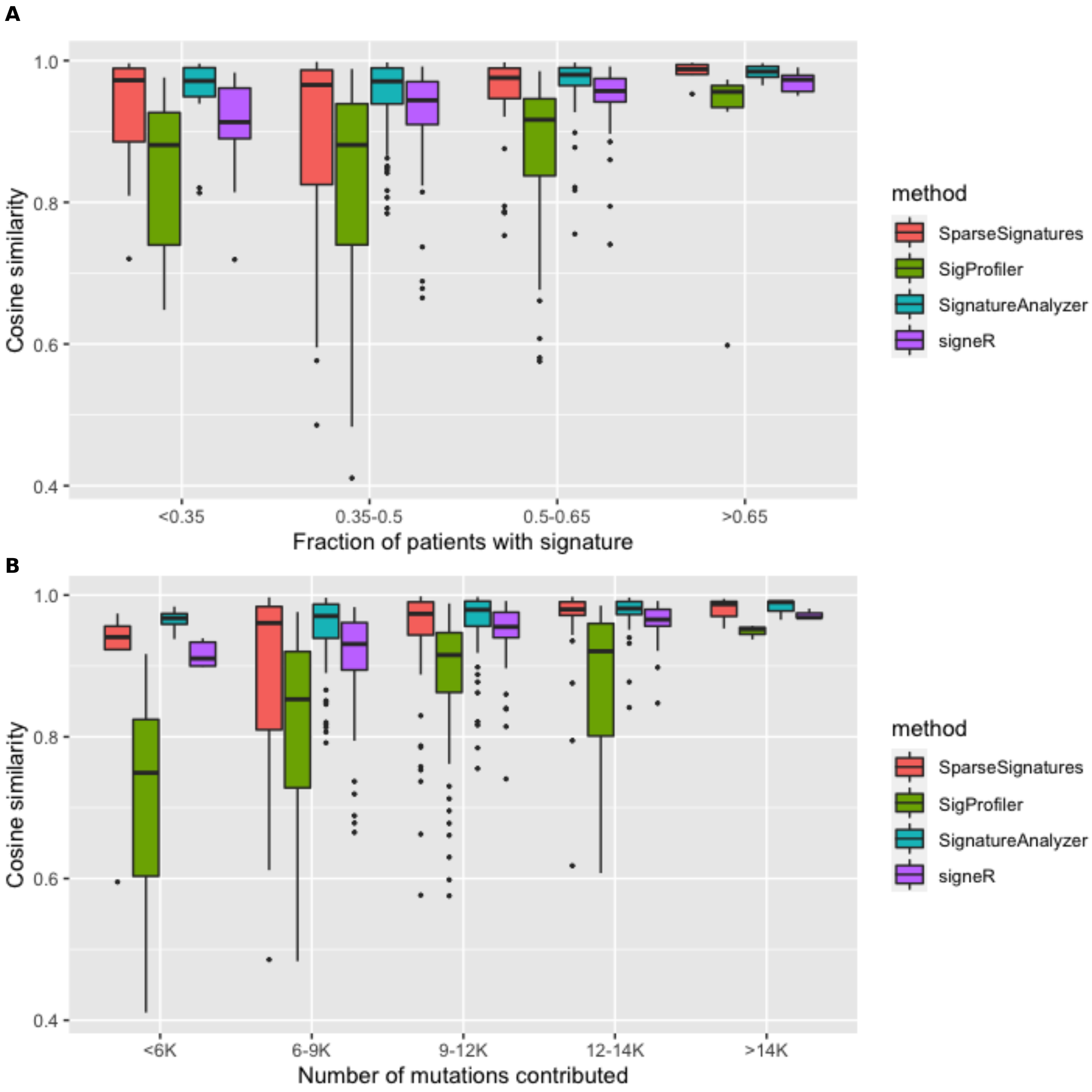

### SupplementaryFigure10.pdf

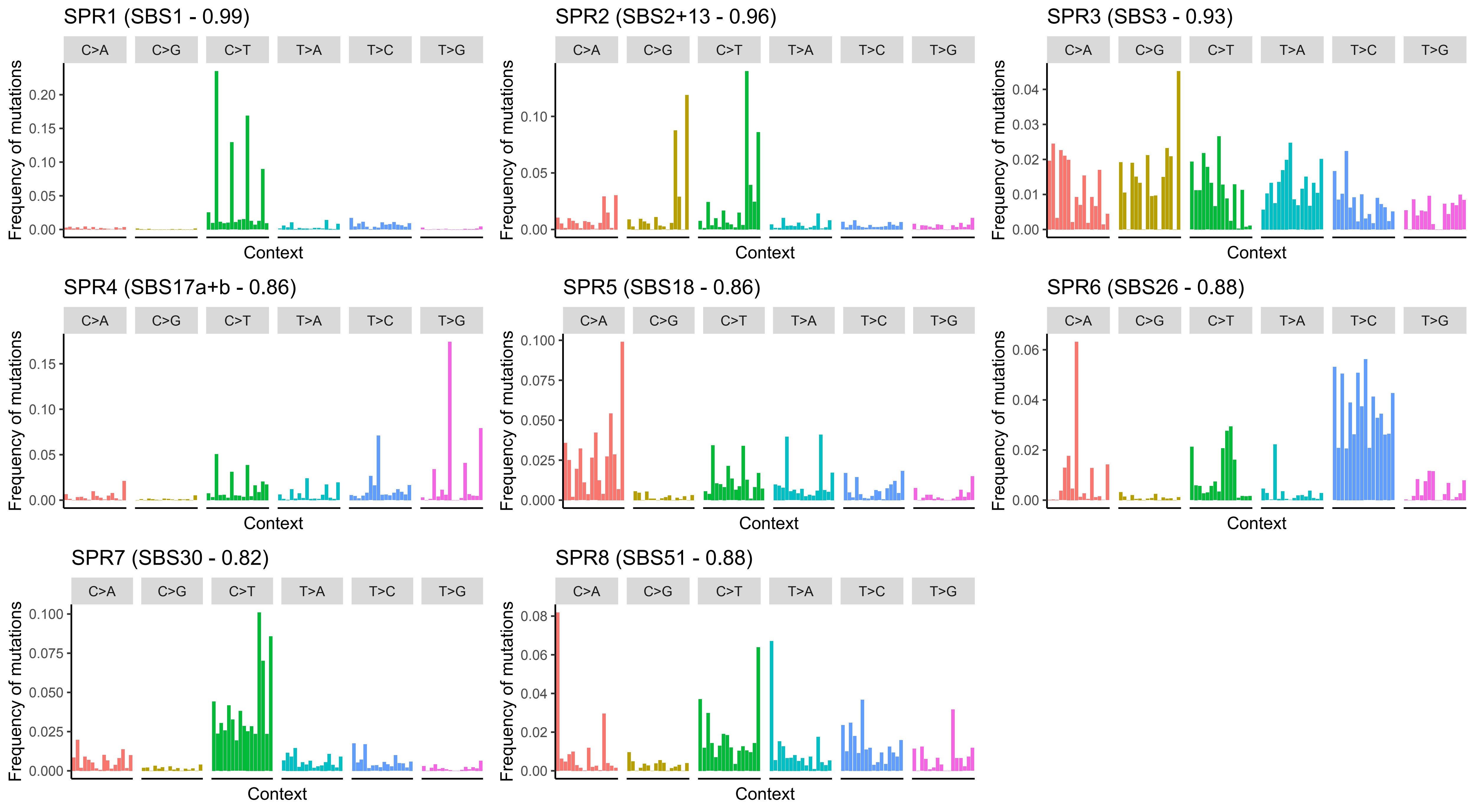

### SupplementaryFigure11.pdf

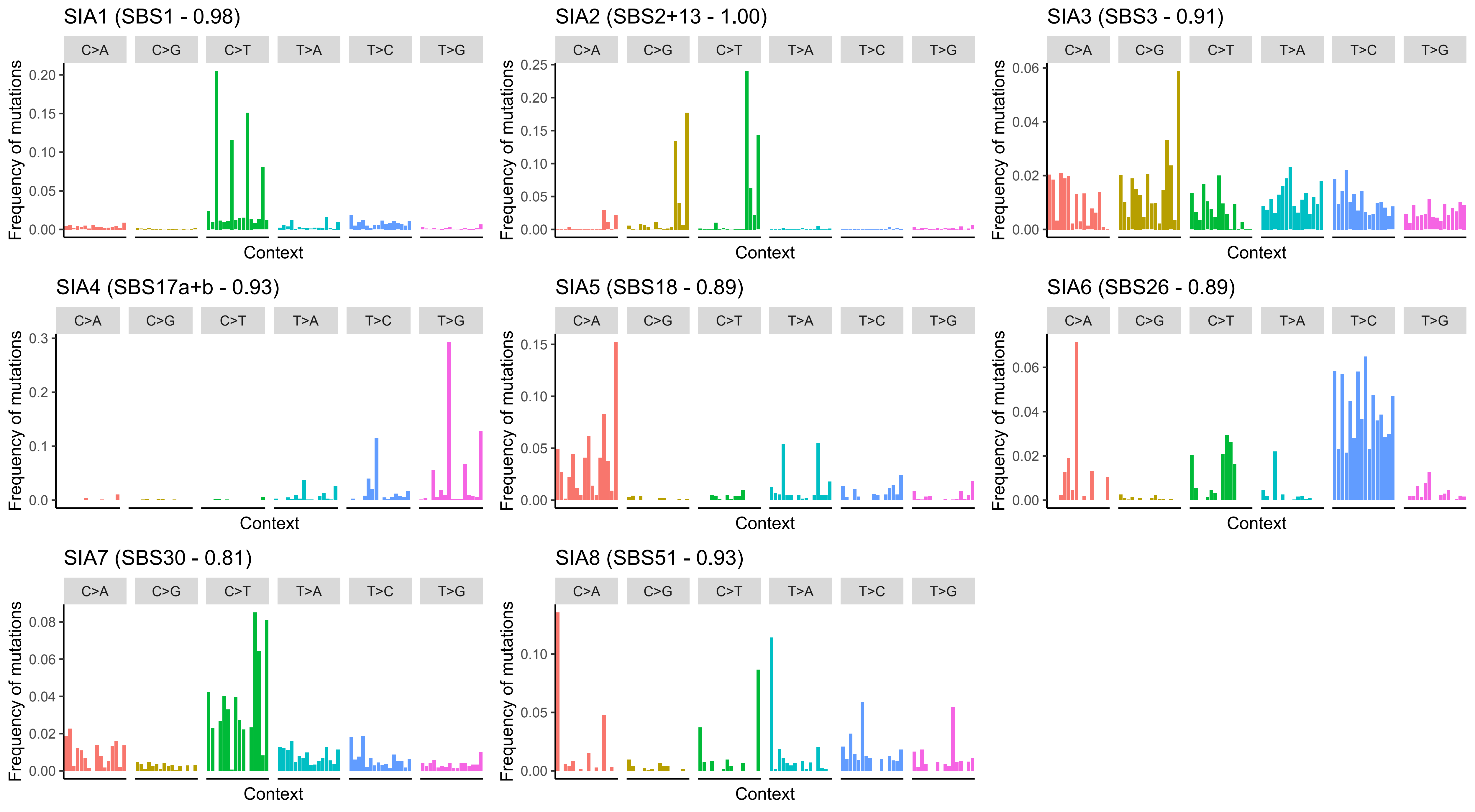

### SupplementaryFigure12.pdf

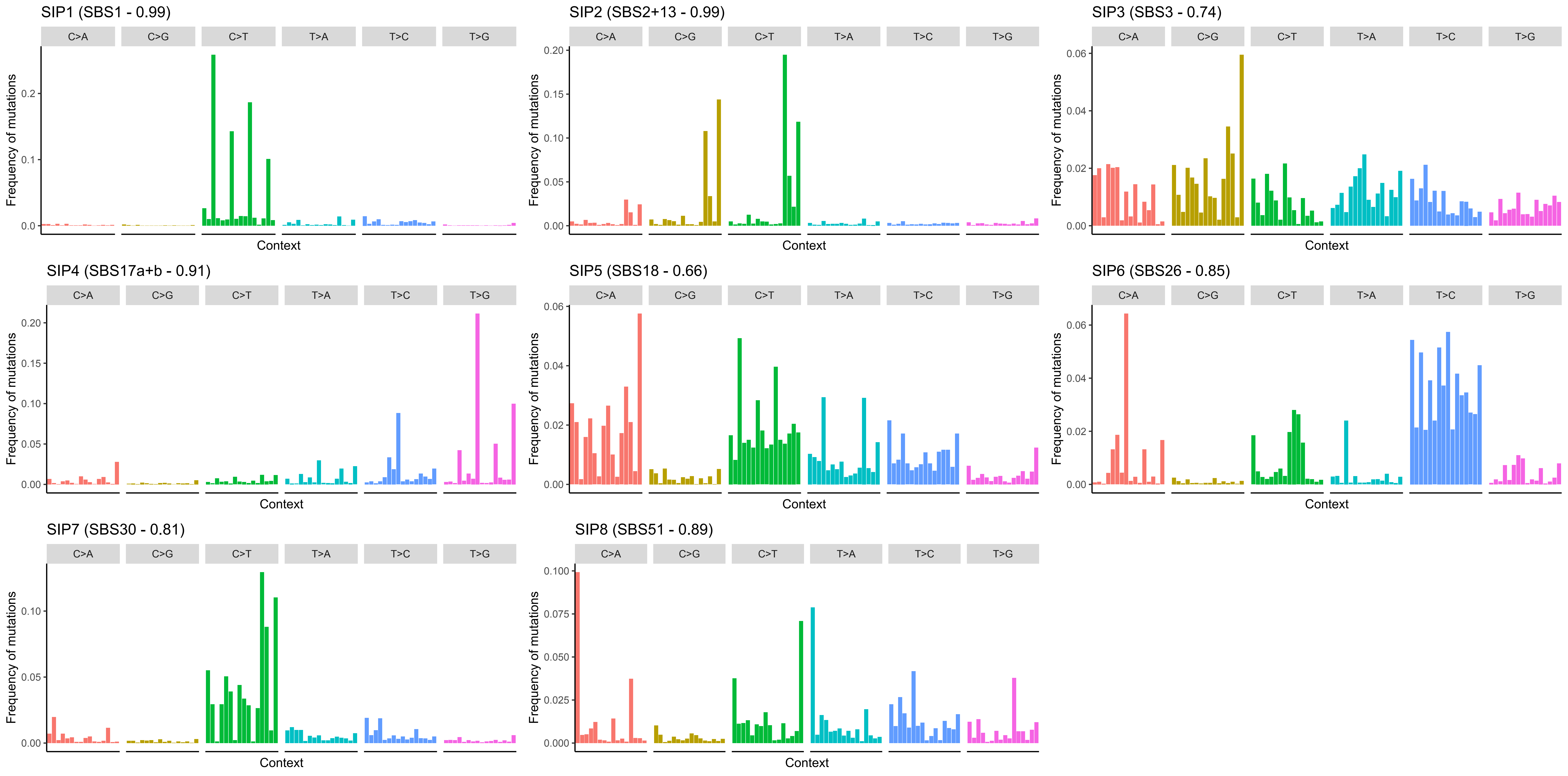

### SupplementaryFigure13.pdf

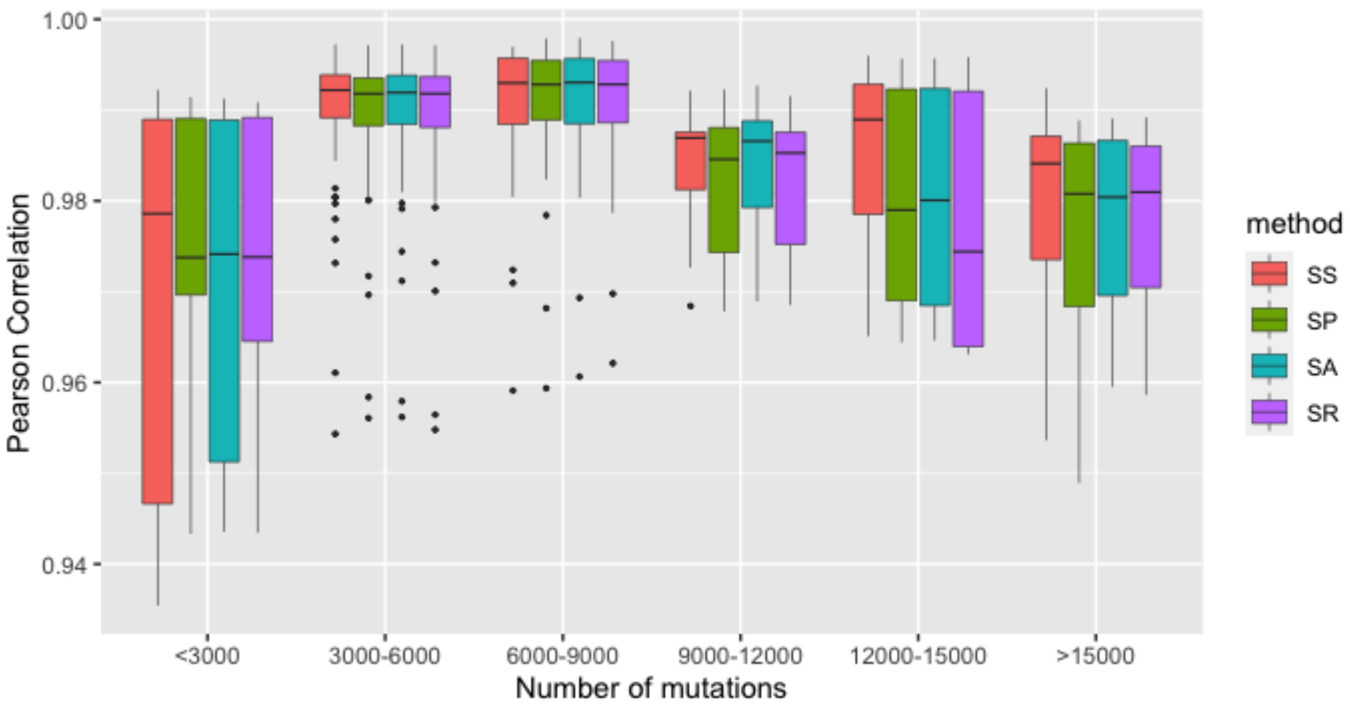

### SupplementaryFigure14.pdf

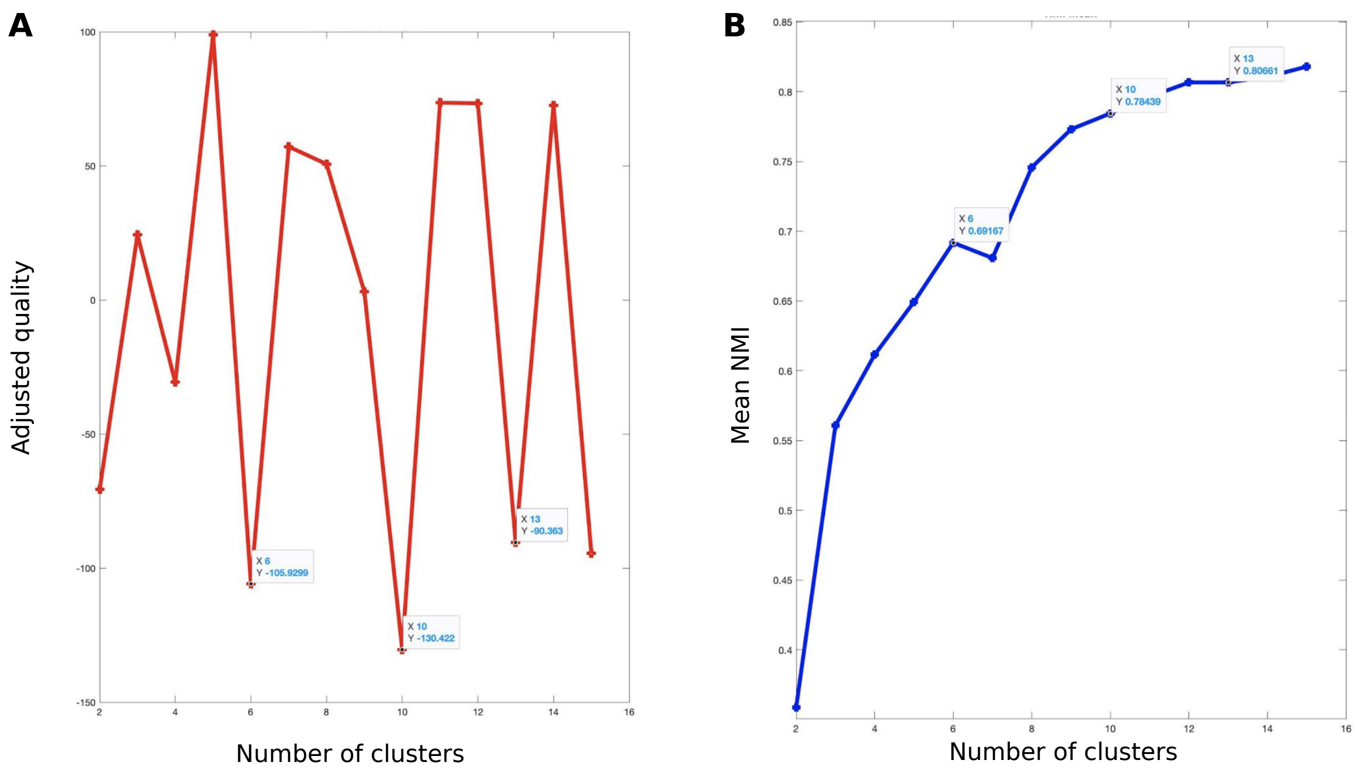

### SupplementaryFigure15.pdf

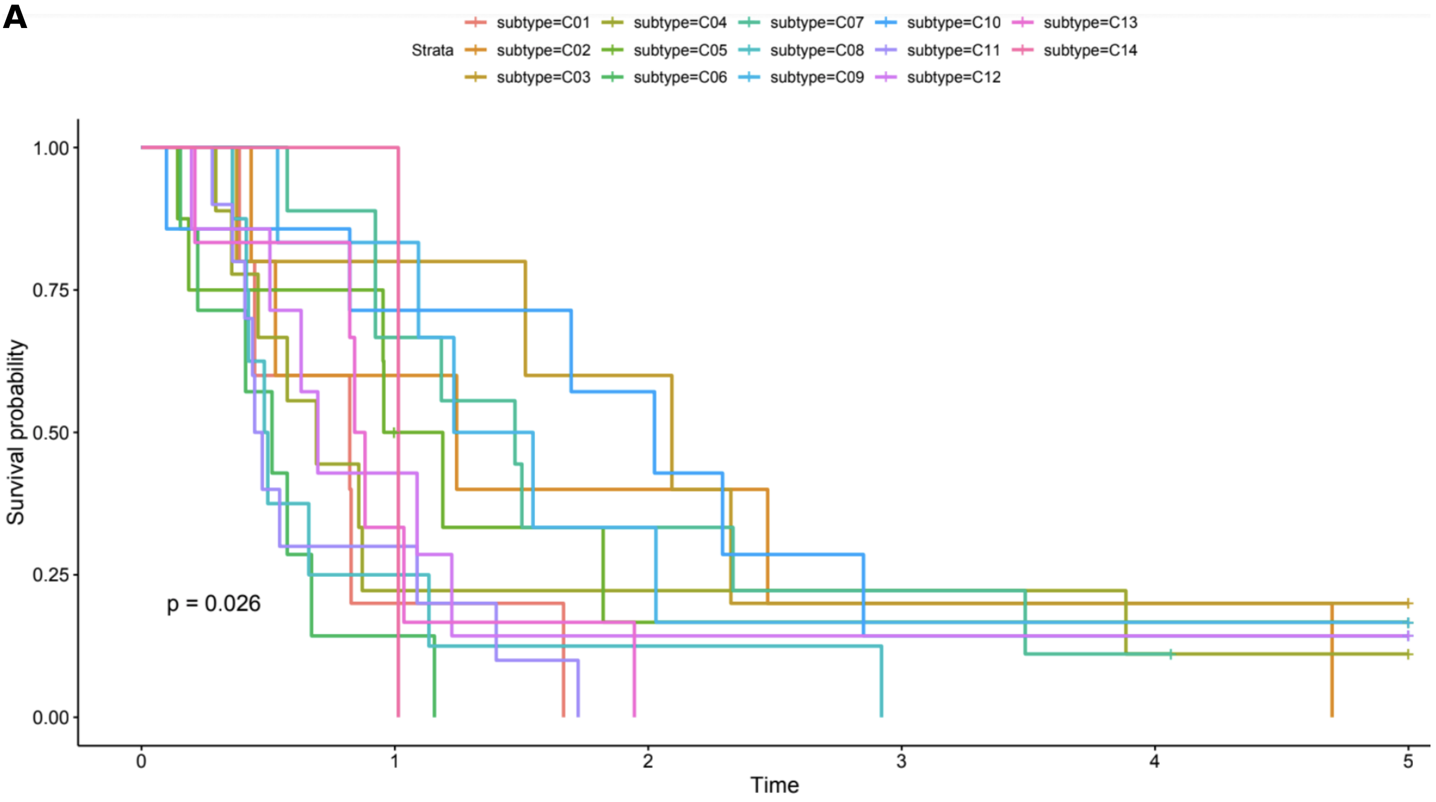

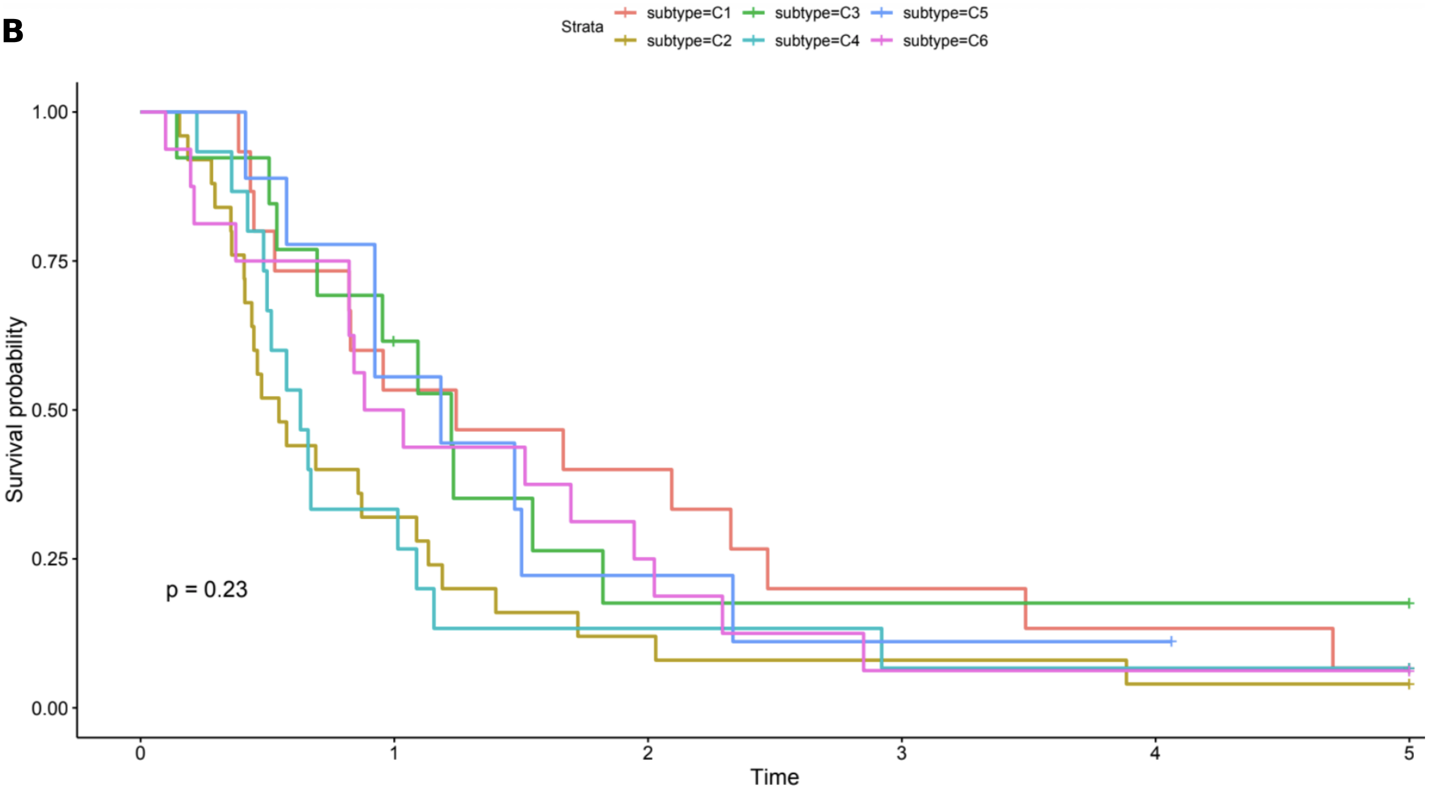

### SupplementaryFigure16.pdf

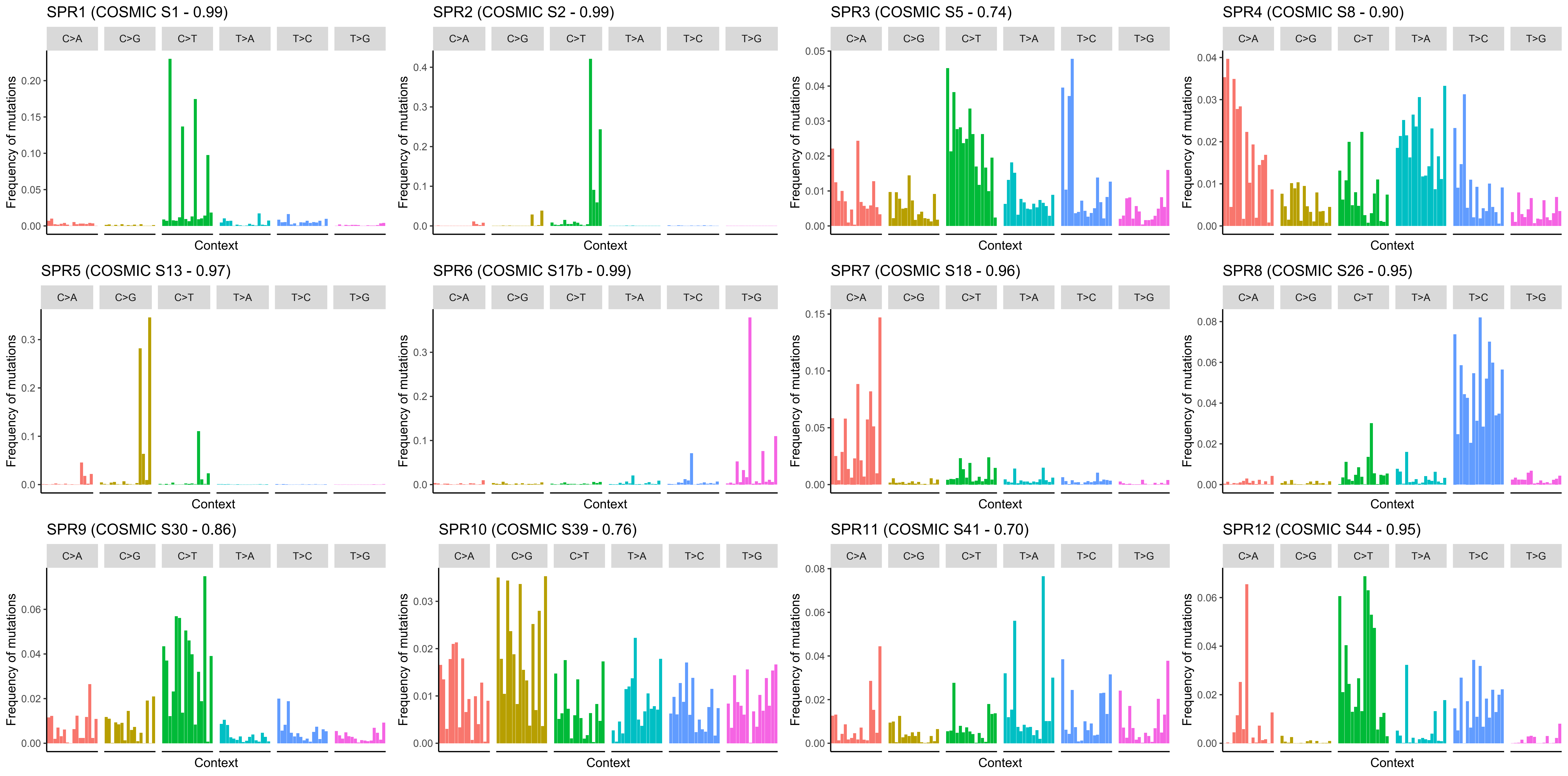

### SupplementaryFigure17.pdf

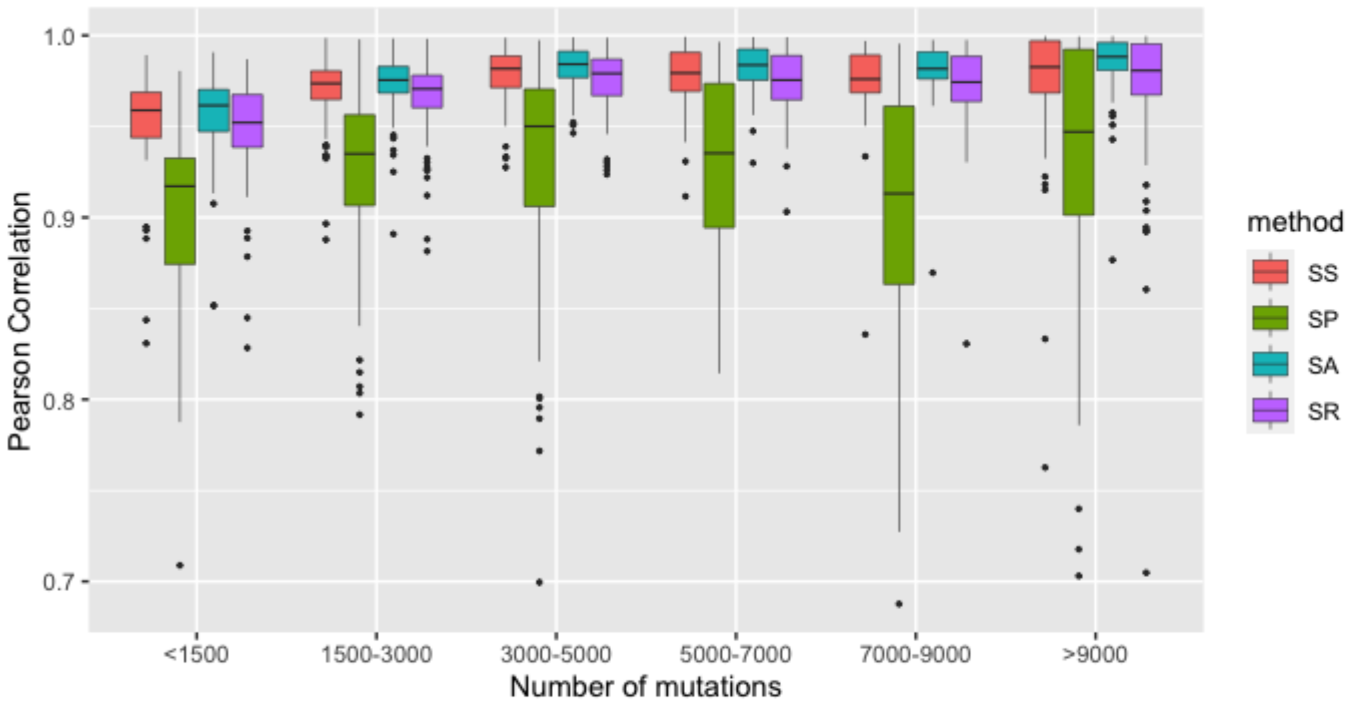

### SupplementaryFigure18.pdf

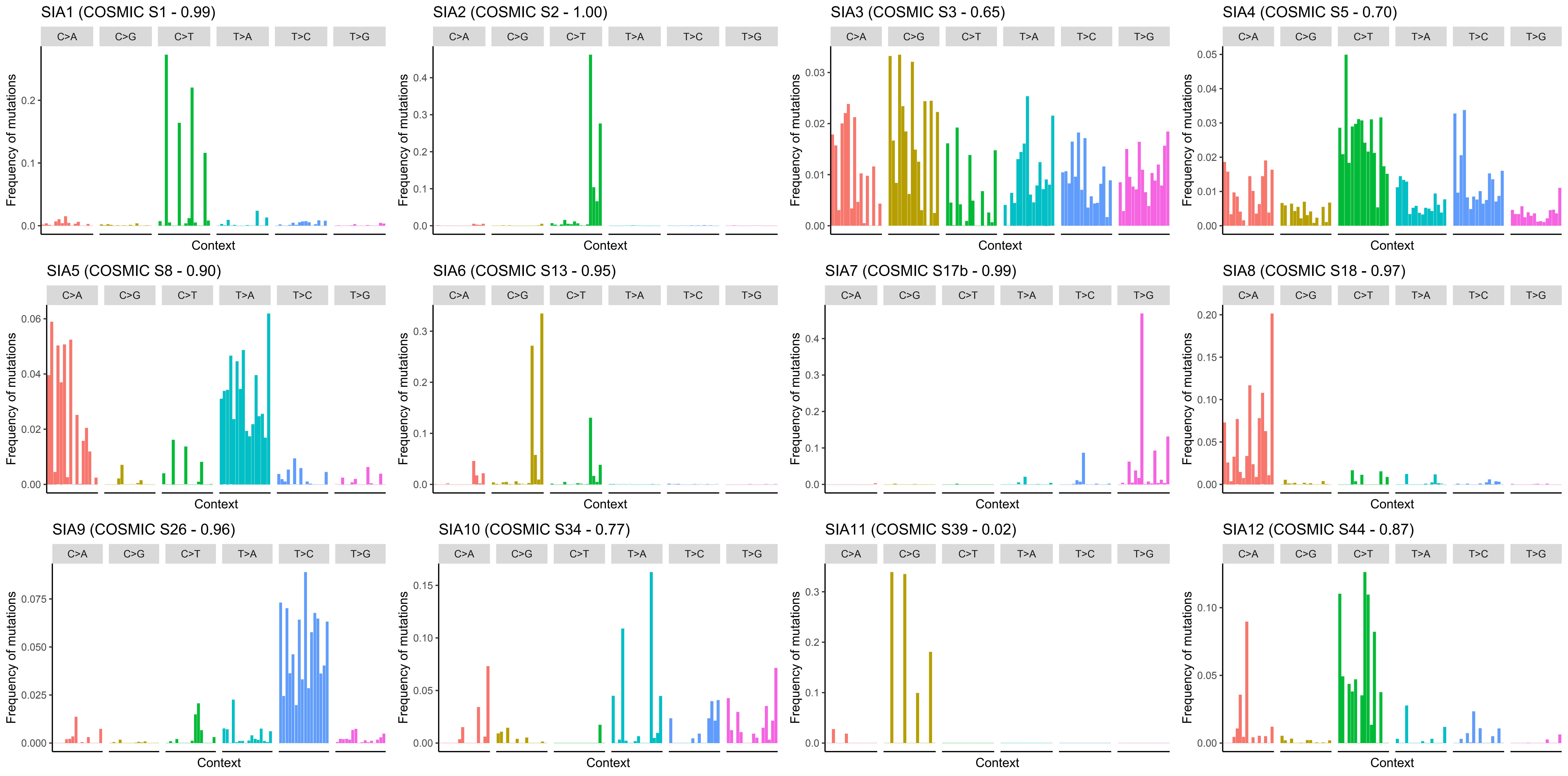

### SupplementaryFigure19.pdf

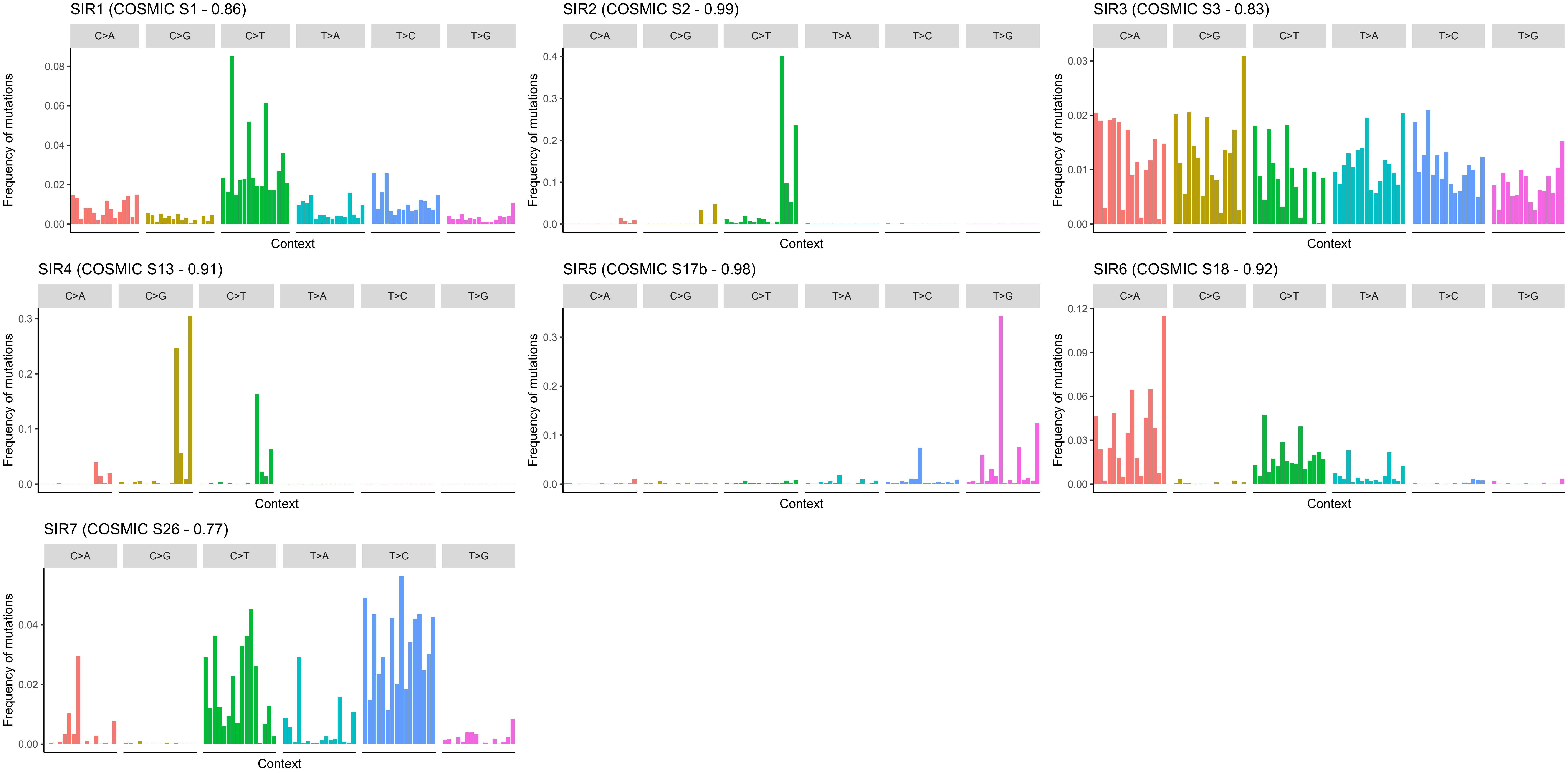
