## Supplementary Methods for "De Novo Mutational Signature Discovery in Tumor Genomes using SparseSignatures"

### 1. Validating the germline-based background signature

We obtained whole-genome sequencing data for 10 randomly selected normal breast tissue samples from Nik-Zainal et al.[1] and ran Sentieon Genomics[2] (default parameters, minimum quality of 30 for calls) on the BAM files to obtain variant calls. We compiled a list of unique high-confidence SNPs across all 10 samples. For all positions where the ancestral allele (obtained from Phase 3 data from the 1000 Genomes project) differed from the reference allele, the ancestral allele was taken in place of the reference allele. We then counted the number of SNPs in each of the 96 trinucleotide-based categories and divided them by the total number of SNPs to obtain the germline mutation spectrum. We found a cosine similarity of 0.98 between this background and the one based on Rahbari et al[3]. The final values used for the germline-based background model in SparseSignatures were the ones from Rahbari et al[3].

### 2. Simulations to choose the percentage of cells in the matrix M to be held out for cross-validation

We selected a set of 10 mutational signatures from the ones listed in COSMIC (https://cancer.sanger.ac.uk/cosmic/signatures_v2) with a known etiology. These are: Signatures 1, 2, 7, 10, 11, 13, 15, 22, 24, 26 in COSMIC.

We then generated random configurations of signatures by selecting the germline background, the methylation signature (Signature 1 from the list above) and 4 additional signatures randomly chosen among the 9 remaining signatures in the list.

For each selected signature, we randomly generated the magnitude of its exposure per patient, by sampling the number of point mutations per signature using a negative binomial distribution with mean 6000 and dispersion parameter 1.5, with the constraints of a minimum number of 1000 mutations per tumor and a maximum of 20000. These parameters were estimated from real data[1]. For each configuration, we generated simulated data from 500 and 1000 tumors.

This procedure was repeated to obtain 100 simulated datasets with 500 tumors and 100 simulated datasets with 1000 tumors.

For cross-validation, values of K ranging from 3 to 7 were tested. $\lambda$ was set to be equal to 0.05.

For each dataset, the initial values of the signature matrix β were computed by 5 repetitions of NMF. Cross-validation was performed by holding out (replacing with 0) 0.1%, 1% and 10% of the entries in the mutation count matrix M. Each cross-validation was repeated 5 times with 5 restarts each.

In these simulations, using 1% of the entries for cross-validation resulted in the most accurate prediction of K, both in datasets containing 500 samples and 1000 samples. Detailed results are reported in Supplementary Table 29.

### 3. Simulations to compare performance of SparseSignatures, SignatureAnalyzer and SigProfiler

We performed 6 independent experiments.

*Simulation 1 (presented in the main text).* We generated simulated data directly by sampling results on real data.

First, we considered a dataset comprising 116 high quality whole genome mutational profiles from the ICGC study with ID *PRAD-CA* (Prostate cancer samples, also included in the curated set by the pan-cancer analysis of whole genomes, PCAWG).

Signatures were assigned to the 116 patients using deconstructSigs[4]; 4 signatures were considered: our background signature and 3 additional signatures which have been observed to be prevalent in prostate cancers (COSMIC SBS1, SBS3, SBS18). All signatures were normalized to sum to 1.

We then simulated the mutation counts in each of the 96 categories for each patient by sampling the chosen number of mutations from each sample and randomly generating noise by adding a value uniformly chosen between 0 and +25 to each of the 96 mutation categories. Mutation counts were generated for 116 patients in this way.

This process was repeated 50 times to generate 50 independent simulated datasets. The methods were executed with the following settings:

SparseSignatures:

K (number of signatures): 2 to 6

$\lambda$ (sparsity): 0.000, 0.025 and 0.050
Number of repetitions of NMF to calculate initial values: 10
Number of iterations to fit signatures using the alternating method with sparsity: 30
Number of repetitions of bi-cross-validation: 10
Number of restarts per repetition of cross-validation: 5

SignatureAnalyzer:

Maximum value of K: 6
Repetitions of complete pipeline: 10

SigProfiler:

Maximum value of K: 6
Repetitions of complete pipeline (input matrix is bootstrapped): 10

signeR:

K (number of signatures): 2 to 6
Repetitions of complete pipeline with defaults: 10

*Simulation 2:* We generated simulated data as for Simulation 1 with the only difference being in how we generated noise; in this case, we added a randomly chosen value between -15 and +15.

*Simulation 3 (signatures randomly selected from COSMIC version 3).*

First, 4 signatures were generated. One signature was fixed - our background signature. The remaining 3 signatures were randomly selected from the whole set of COSMIC signatures version 3 (removing SBS5, which has a 0.98 similarity with our background signature). All signatures were normalized to sum to 1.

We then generated the number of mutations from each signature for 100 patients. For the background signature, this was done by sampling from a negative binomial distribution with parameters (size = 1.5, mean = 2000). This signature was therefore present in all samples. For the other signatures, we first randomly selected 33-67 patients in whom the signature was present. For these patients, the number of mutations from each randomly generated signature was obtained by sampling from a negative binomial distribution with parameters (size = 1.5, mean = 200).

We then simulated the mutation counts in each of the 96 categories for each patient by sampling the chosen number of mutations from each signature. Finally, randomly generated noise was included by adding a value uniformly chosen between 0 and +25 to each of the 96 mutation categories. Mutation counts were generated for 100 patients in this way.

This process was repeated 50 times to generate 50 independent simulated datasets. The inference methods were executed with the same settings as for Experiment 1.

*Simulation 4 (signatures randomly selected from COSMIC version 3 dense signatures).*

We generated simulated data as for *Simulation 3* with the only difference being in how we selected the signatures at the beginning of the simulations. In this case, we select our background plus 3 additional signatures from COSMIC version 3, only considering dense signatures. In particular, we considered only COSMIC signatures with <0.001 contribution for <25% of the 96 trinucleotides, i.e., these signatures present high contribution for most of the 96 trinucleotides. Specifically, we considered the following COSMIC version 3 signatures: SBS3, SBS4, SBS8, SBS9, SBS16, SBS18, SBS24, SBS25, SBS31, SBS35, SBS36, SBS37, SBS38, SBS39, SBS40, SBS41, SBS42.

*Simulation 5 (signatures randomly selected from COSMIC version 3 sparse signatures).*

We generated simulated data as for *Simulation 3* with the only difference being in how we selected the signatures at the beginning of the simulations. In this case, we select our background plus 3 additional signatures from COSMIC version 3, only considering dense signatures. In particular, we considered only COSMIC signatures with <0.001 contribution for >50% of the 96 trinucleotides, i.e., these signatures present a low contribution for most of the 96 trinucleotides. Specifically, we considered the following COSMIC version 3 signatures: SBS1, SBS2, SBS6, SBS7a, SBS7b, SBS10a, SBS10b. SBS11, SBS13, SBS15, SBS17a, SBS17b, SBS21, SBS23, SBS28, SBS30, SBS34.

*Simulation 6:* We generated simulated data as for Experiment 3 with the only difference being in the number of selected signatures. In this case, each simulated dataset includes 8 signatures, including the background signature plus 7 additional signatures randomly chosen from COSMIC version 3.

**4. Methods and parameters used for signature discovery on real data**

SparseSignatures:

K (number of signatures): 2 to 18
λ (sparsity): 0.01, 0.05, 0.10, 0.20
Number of repetitions of NMF to calculate initial values: 200
Number of iterations to fit signatures using the alternating method with sparsity: 30
Number of repetitions of bi-cross-validation: 200
Number of restarts per repetition of cross-validation: 5

SignatureAnalyzer:

Maximum value of K: 18
Repetitions of complete pipeline: 200

SigProfiler:

Maximum value of K: 18
Replicates per iteration: 200

signeR:

K (number of signatures): 2 to 18
Repetitions of complete pipeline with defaults: 200

**5. Estimation of the optimal number of clusters for CIMLR**

We used two heuristic approaches to estimate the optimal number of clusters for CIMLR. The first approach is described in Ramazzotti et al.[5] and consists in evaluating the eigengap of each low-rank approximation; considering this heuristic, the optimal number of clusters is the one where the drop in eigengap is maximum. In addition, we here also adopted a second heuristic aimed at assessing the stability of clustering solutions for the various low-rank approximations. Specifically, we first computed the clustering assignments for number of clusters varying from 2 to 10 on the complete dataset and then repeated the same procedure on 100 datasets obtained via bootstrap by resampling; stability of clustering solutions was estimated as the mean normalized mutual information across the 100 bootstrap inferences.
